# Imaging-based spatial isoform transcriptomics resolves transcript diversity at cellular and subcellular resolution

**DOI:** 10.64898/2026.09.22.753429

**Authors:** McAndrew Eamon, Fierville Morgane, Gautier-Isola Marine, Pignol Marie, Arguel Marie-Jeanne, Cadis Hugo, Barrès Romain, Mari Bernard, Barbry Pascal, Lebrigand Kevin

**Author notes:** Contributed equally (McAndrew Eamon, Fierville Morgane). Correspondence to Kevin Lebrigand, Institut de Pharmacologie Moléculaire et Cellulaire, 660 route des Lucioles, 06560 Valbonne.

## Abstract

Spatial transcriptomics has transformed our understanding of tissue organization, yet current high-throughput approaches remain largely confined to a gene-centric paradigm, precluding systematic characterization of spatially resolved transcript isoform diversity. Here, we introduce imaging-based spatial isoform transcriptomics (iSiT), a framework enabling simultaneous large-scale gene expression and targeted transcript isoform profiling on the commercial Xenium platform. iSiT resolves alternative splicing events that alter internal transcript structure at cellular and subcellular resolution using a single junction-specific probe per target. To facilitate broader adoption, we developed ProbeHarbour, an open-source toolkit for the design, characterization and management of isoform-specific probes. We demonstrate iSiT in the postnatal mouse brain by combining the Prime 5K gene-level panel with isoform-specific detection of 45 synaptogenesis-associated genes. By extending spatial transcriptomics beyond gene-level measurements, iSiT provides a scalable and readily accessible framework for investigating spatial isoform regulation in development and disease.

## Introduction

In recent years, spatial transcriptomics (ST) has transformed our ability to investigate tissue organization by enabling the mapping of gene expression within intact biological contexts^1,2,3^. Combined with single-cell RNA sequencing (scRNA-seq), ST has provided insight into cellular heterogeneity, developmental trajectories and tissue architecture across diverse biological systems, including the nervous system^4,5^. Alternative splicing is a major source of transcript and protein diversity, generating isoforms with distinct molecular properties, including altered protein domains, interaction partners, subcellular localization and regulatory functions^6,7^. However, current high-throughput ST platforms remain largely confined to a gene-centric paradigm, limiting the systematic characterization of transcript isoform diversity within tissues^8^. Resolving transcript isoforms in their native spatial context is therefore an essential next step toward a more complete understanding of cellular states and tissue function.

Recent advances have begun to address the limitations of gene-level transcriptomics by integrating long-read sequencing with scRNA-seq or ST approaches. Long-read single-cell methods, including ScNaUMI-seq^9^, ScISOr-Seq^10^, and LR-Split-seq^11^, have enabled comprehensive characterization of isoform diversity within complex cell populations. Similarly, spatial isoform profiling approaches have combined spatial barcoding with long-read sequencing to extend isoform analysis into intact tissues. SiT^12^, for example, integrates long-read sequencing with the 10x Genomics Visium platform to enable spatial transcript isoform detection. However, spots of 55 µm typically capture transcripts from multiple cells lacking true single-cell resolution. More recent approaches, including Spl-ISO-Seq^13^ and Spl-ISO-Seq2^14^, have improved spatial resolution leveraging the Slide-seqv2 and Stereo-seq ST platforms. Nevertheless, these approaches remain constrained by limited capture sensitivity and high dropout rates. Because they rely on 3′ capture and template switching, recovery of transcripts longer than 2 kb is inefficient, giving incomplete representation of transcript diversity^15^.

Imaging-based approaches provide an alternative by directly detecting transcripts within their native environments. A MERFISH-based platform, RT&T-AMP-MERFISH^16^ enabled spatial profiling of ∼10,000 transcript isoforms in the mouse brain, establishing that isoform-resolved ST is feasible at scale. Yet it is a custom implementation rather than a commercially available system, which currently limits its wider adoption. Furthermore, isoform discrimination relied on multiple probes (4-6), requiring a minimum of 120 nt of private sequence per transcript. This limited isoform profiling largely to transcript-end variation, such as 3′ and 5′ untranslated region usage. Therefore several important classes of alternative splicing events, including cassette exon inclusion and exclusion, mutually exclusive exons and cryptic exon usage, remain difficult to interrogate.

Alternative splicing plays a particularly prominent role in the nervous system^17^, where extensive transcript diversity contributes to neuronal identity, synaptic specificity and circuit plasticity^18^. Developmentally regulated cassette exons constitute one of the predominant mechanisms shaping transcriptomic diversity during brain development^19^. Thus, a scalable and broadly applicable framework capable of resolving diverse isoforms arising from internal transcript structure while maintaining compatibility with broadly accessible high-throughput spatial transcriptomics platforms remains lacking.

Here, we introduce imaging-based spatial isoform transcriptomics (iSiT), a framework that extends the capabilities of commercial Xenium ST toward isoform-resolved analysis. By combining large-scale gene expression profiling with targeted isoform detection, iSiT enables spatial characterization of transcript isoforms and resolves common classes of alternative splicing events at cellular and subcellular resolution, using as few as one probe per target. We applied iSiT to the developing mouse brain, where extensive transcript diversity contributes to the formation and maturation of neural circuits^20,21^. In mammals, synaptogenesis rapidly accelerates during the early postnatal period^22^, particularly in brain regions such as the cortex and hippocampus. This process depends on the precise spatiotemporal regulation of molecular programs, many of which are shaped by alternative splicing^23^. Disruptions in synaptogenesis have been implicated in neurodevelopmental disorders, including autism spectrum disorder, schizophrenia and intellectual disability^24,25^. Unraveling the molecular underpinnings of synapse formation is not only central to developmental neuroscience but also critical for identifying the origins of these conditions. Across six postnatal developmental stages, iSiT revealed spatially and temporally regulated isoform programs involving key regulators of synaptic adhesion, neurotransmission and vesicle trafficking, including *Clta*, *App*, AMPA receptor subunits and neurexins. Our study provides a scalable framework for investigating transcript isoform regulation across development, physiology and disease at cellular and subcellular resolution.

## Results

### A spatiotemporal atlas of the developing mouse brain

To systematically map the developing mouse brain cellular landscape, we employed an in situ imaging-based assay using the 10x Genomics Xenium platform. The platform detects transcripts in place using padlock probes, which require adjacent hybridization of both arms before ligation, followed by rolling-circle amplification and sequential imaging of individual transcript molecules at subcellular resolution (∼100 nm). We used the Prime 5K chemistry, allowing single-molecule quantification of 5,006 genes. Experiments were conducted across six early postnatal stages (P1, P3, P5, P7, P14, and P21), with two biological replicates per stage, one male and one female, and a single hemisphere analysed per animal (**Fig.1a**). In total, our dataset comprises 1,348,351 cells (**Fig.1b**), with a mean of 1,124 transcripts detected per cell (**Fig.1c**) and more than 2.2 billion high-quality transcripts (QV≥20) (**Fig.1d**). Following dataset integration using Harmony^26^ and unsupervised Leiden clustering, cell identities were assigned by label transfer based on transcriptomic similarity to an adolescent mouse brain (P12-P30) reference single-cell atlas^4^. To ensure robust annotation across developmental stages, we anchored labelling to Harmony-integrated Leiden clusters, incorporating gene markers, spatial context, and label-transfer information yielding 31 major cell types (**Fig.1e-f**, **,Extended Data Fig. 1** and **Methods**). Left and right hemispheres showed concordant cell-type composition, indicating that analysing a single hemisphere introduces no systematic bias (**Extended Data Fig. 2a,b**), and male and female animals showed concordant composition, supporting their use as independent biological replicates within each stage (**Extended Data Fig. 2c,d**). Cell-type proportions changed markedly over time, with progressive declines in neuroblasts, glia-like, Cajal-Retzius, peptidergic, cholinergic and monoaminergic populations and expansion of mature types including cortical layer (CTX) and dentate gyrus (DG) excitatory neurons, astrocytes, microglia and oligodendrocytes. Oligodendrocyte maturation proceeds from precursors (OPC) through committed progenitors (COP) to myelinating cells (MFOL) (**Fig.1g-h**).

**Fig. 1:**
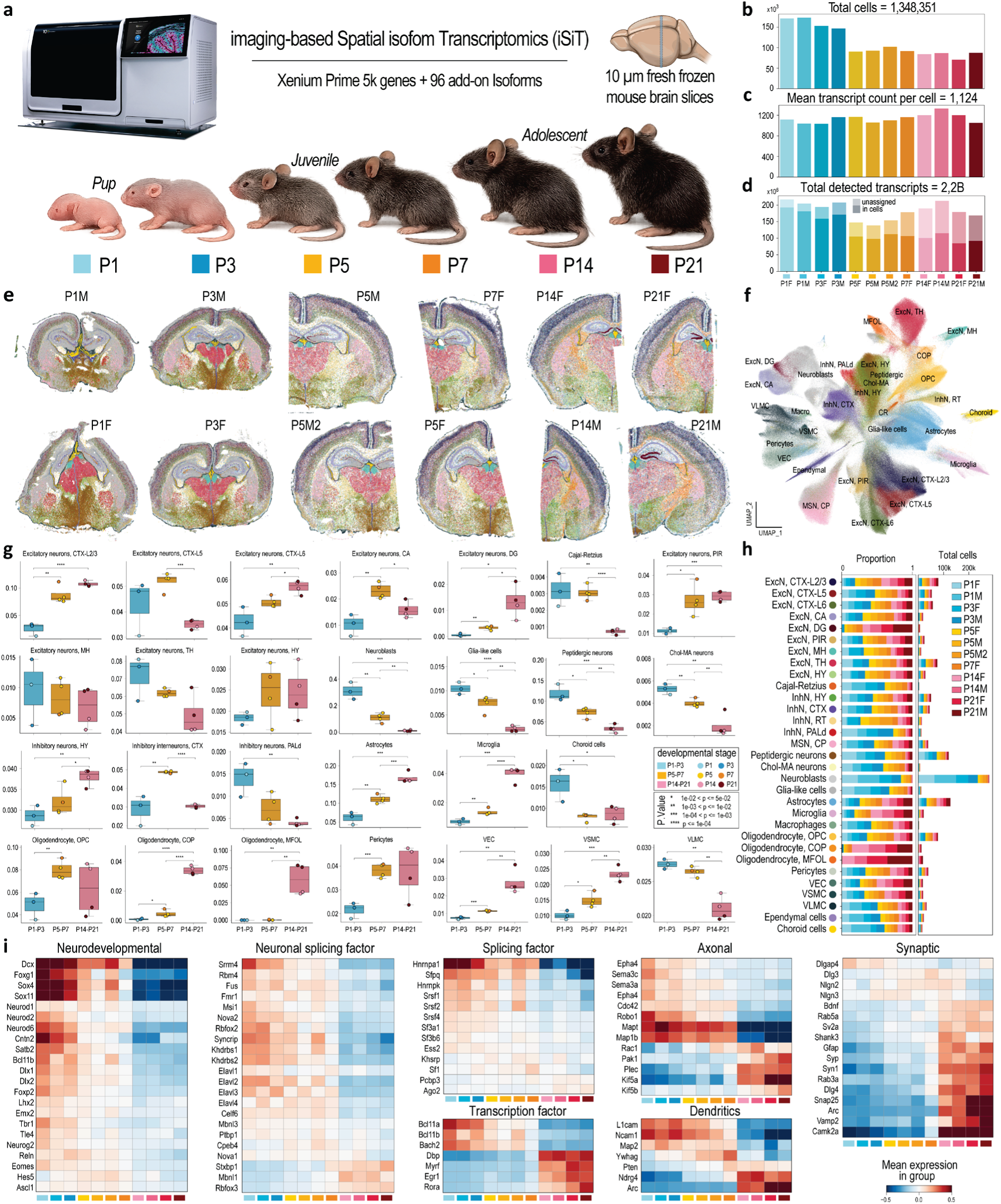
Spatial Transcriptomics of the developing mouse brain (P1-P21). **a**, iSiT experimental design. **b**, Total number of cells per sample. **c**, Mean detected transcripts per cell. **d**, Total inside (solid) and outside (shaded) cell segmentation mask-detected transcripts per sample. **e**, Spatial map colored by cell type. **f**, Harmony-integrated UMAP colored by cell type. **g**, Box plots showing cell type proportions across hemispheres, grouped into three stages of development (P1-P3, P5-P7 and P14-P21); dots colored by individual time points from P1 to P21. **h**, Cell type proportions (left) and total cell numbers (right) across hemispheres, colored by developmental stage. (**i**) Scaled expression of representative genes involved in neurodevelopmental processes, transcriptional regulation, neuronal identity, and RNA splicing, together with axonal, dendritic, and synaptic compartment markers, across mouse brain maturation.

We next examined the temporal expression of key gene groups, including neurodevelopmental genes, splicing factors, transcription regulators and established markers of axons, dendrites and synapses (**Fig.1i**). Neurodevelopmental and splicing regulator expression declined progressively from P1 to P21, whereas synaptic and dendritic genes showed a pronounced increase from P14. Gene-level analysis cannot resolve the transcript isoform diversity generated by alternative splicing. We therefore extended the framework to the isoform level, enabling identification of transcript-specific developmental regulation.

### Cross-platform comparison of isoform-resolved spatial transcriptomics approaches

Isoform-resolved transcriptomics at scale has relied on poly(dT) capture in single-cell and ST protocols coupled with long-read sequencing. We^12^ and others^27^ have previously demonstrated that these technologies can uncover isoform switches that remain obscured by conventional short-read protocols. However, their broader adoption is constrained by low capture efficiency^28^ and by a length bias favoring short transcripts (mean <1 kb)^29^, which together impede robust detection of long transcripts. We compared probe-based (Xenium) and poly(dT)-based (Visium HD; 334M short reads; 84.9% sequencing saturation) capture on adjacent sections of the same sample (P14F). Gene-level expression was well correlated across the 4,954 shared genes (Pearson r = 0.91), but Xenium recovered on average 42-fold more counts per gene than Visium HD nUMI. This gain increased with mean transcript length per gene and with the number of probesets tiling each gene. Within the 1-2 kb transcript length bin, mean gain increased from 30-fold for genes with one probeset (n = 36) to 138-fold for genes with four probesets (n = 8) (**Fig.2a-c**).

To overcome these sequencing-based limitations, we moved to imaging-based spatial transcriptomics on the Xenium platform. Building on the Prime 5K panel, we designed a custom panel of 96 exon-exon junction probes, arranged as mutually exclusive pairs, across 45 genes reported to undergo developmentally regulated isoform switching during brain maturation (**Supplementary Tables 1-3**). Of these 45 genes, 28 were already targeted at the gene-level in the Prime 5K panel. Using a publicly available 10x Genomics Xenium P56 mouse brain section, we demonstrated that integration of the isoform panel preserved gene-level assay performance, with no significant differences in transcript counts or quality values (QV) compared with standard gene-level profiling. (**Fig.2d-g**).

We next benchmarked iSiT against a Visium HD dataset from P56 mouse brain dataset generated using poly(A)-based in situ capture and sequenced across four ONT flowcells (456M long reads; 73.4% sequencing saturation). In total, the long-read dataset assigned 36 million UMIs to an isoform, compared to 1.6 million isoform-resolved transcripts detected in our P14F iSiT sample, which targeted 96 alternative splicing events (**Fig.2h-i**). Restricting the analysis to the 45 genes included in our panel, direct comparison of total isoform-level molecules revealed markedly higher sensitivity with iSiT relative to the long-read approach (mean=37-fold) even for the 28 genes having dedicated gene-level probes on the Prime 5K panel (black-dash borders) and suffering from gene-level probeset competition to access RNA molecules (**Fig.2j**).

Despite its transcriptome-wide scope, the long-read strategy exhibited high sparsity, reflecting intrinsic limitations in capture efficiency and transcript representation. These constraints compromise the quantitative accuracy and resolution of isoform-level measurements, ultimately limiting the capacity of poly(A)-based long-read profiling to finely resolve the biological complexity of alternative splicing in situ, as illustrated by representative examples discussed in the following sections. While short transcripts such as *Clta* (1.1kb) are well quantified, long multi-exonic genes suffer from incomplete isoform coverage, reducing reliability. This affects genes like AMPA receptor subunits (>4kb), leading to limited assignment and, in the worst-case scenario, misassignment of abundance, for example, overestimating non-neuronal App751 over the neuronal App695 isoform (**Extended Data Fig. 3**).

**Fig. 2:**
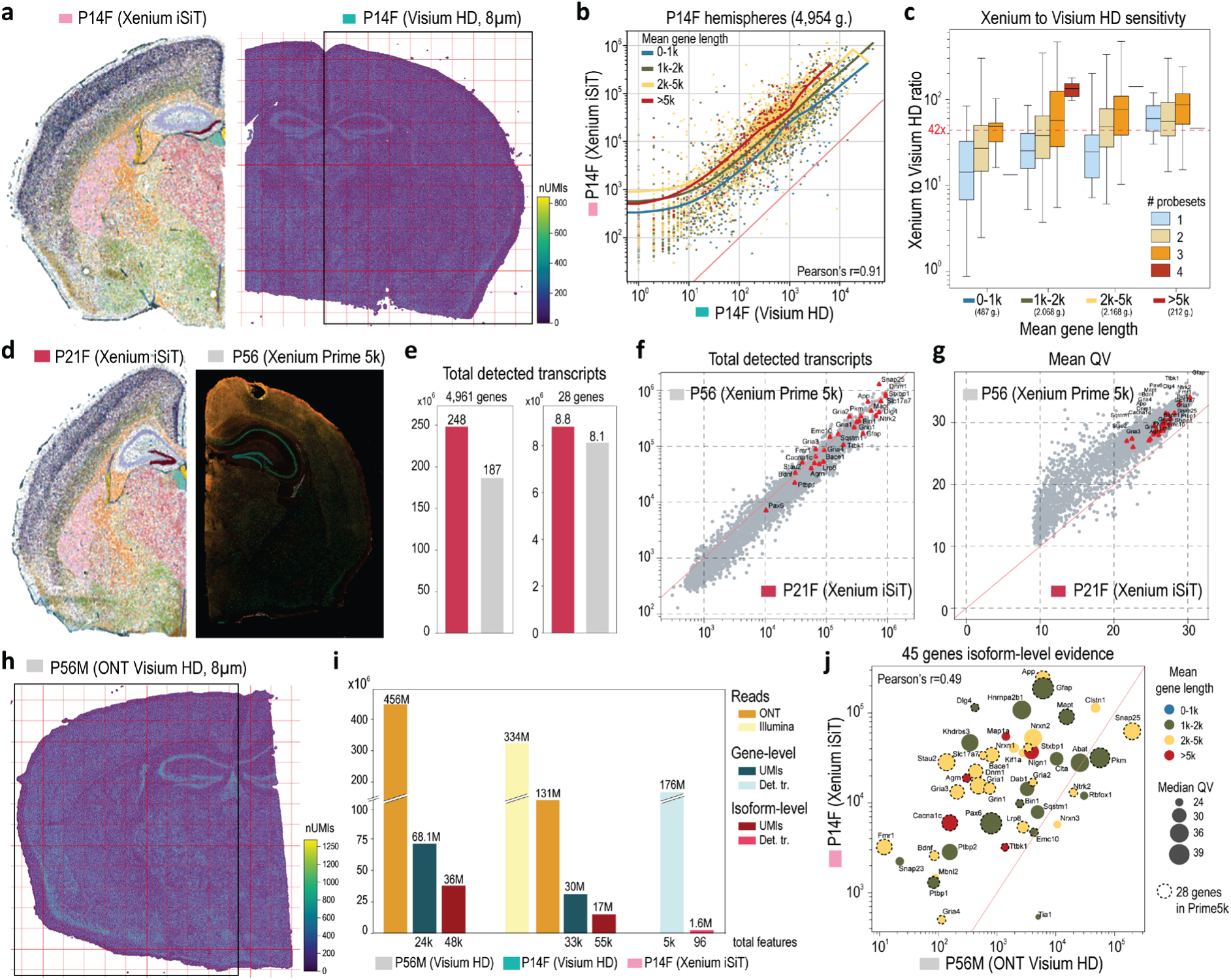
Isoform-level Spatial Transcriptomics: comparison of analytical sensitivity. **a**, Comparison of the P14F sample with a 3′ poly(A)-based in situ capture Visium HD assay. The representative hemisphere region analyzed by Visium HD is indicated by a black rectangle. **b**, Gene-level correlation between P14F hemispheres across 4,954 commonly expressed genes, colored by mean gene length. Loess curves for four gene-length categories show that Xenium sensitivity increases relative to Visium HD with increasing gene length. **c**, Box plots showing the ratio of Xenium-detected transcripts to Visium HD nUMIs, stratified by gene-length categories and by the number of probe sets in the Xenium Prime 5K assay. Total gene number per length category is indicated in parenthesis. **d**, Comparison of the P21F sample with an external 10x Genomics Xenium Prime 5K public dataset from a coronal section of a P56 mouse brain. **e**, Total detected transcripts across both hemispheres for 4,961 commonly expressed genes (left) and for the 28 genes included in both gene-level and isoform-level panels. **f**, Total detected transcripts per gene, showing that the Isoform-level panel has no impact on Xenium Prime 5K quantification. **g**, Mean quality value (QV) per gene, indicating that the Isoform-level panel has no impact on Xenium Prime 5K detection. **h**, External Visium HD dataset from a P56 mouse brain coronal section sequenced using Oxford Nanopore Technologies long-read sequencing. **i**, Comparison of statistical metrics across ONT Visium HD, in-house P14F Visium HD (short- and long-read sequencing), and P14F iSiT sample (Prime 5K plus isoform add-on), at both gene- and Isoform-level resolution. **j**, Isoform-level evidence comparison between P56 ONT Visium HD (nUMIs) and our P14F iSiT (detected transcripts) sample grouped by gene for the 45 genes in the Isoform-level add-on panel.

**Fig. 3:**
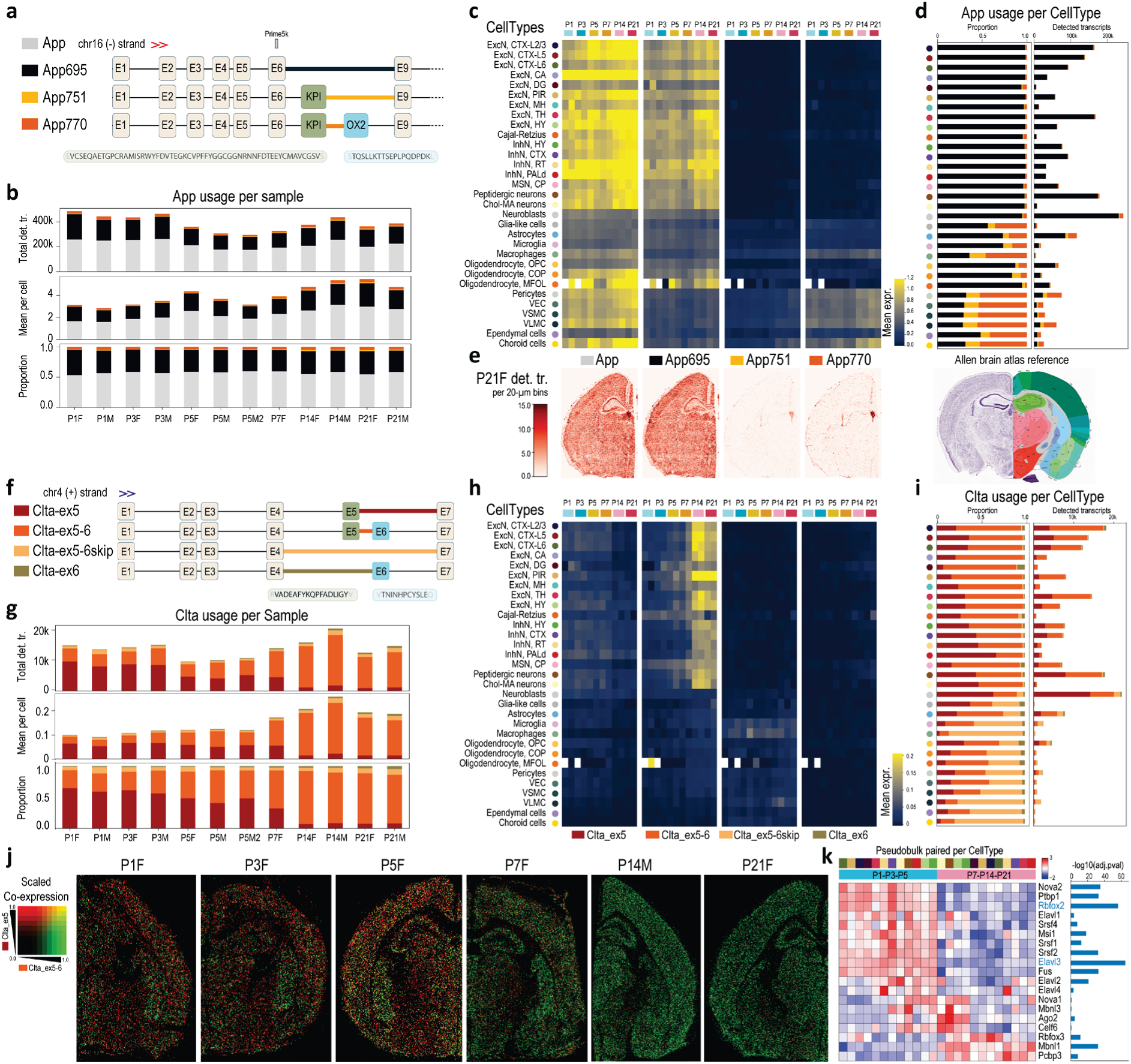
App and *Clta* isoform usage across cell type, space and time. **.a**, *App* gene locus showing three isoforms: App695 lacks exons 7 and 8, App751 includes exon 7 (KPI domain), and App770 includes exons 7 and 8 (KPI+Ox-2 domains). **b**, *App* usage across samples shows a slight increase in the mean number of detected transcripts per cell (middle) and no change in isoform usage across development. **c**, Mean *App* expression per cell type measured by gene-level Prime 5K probeset (left) and isoform-level probes (right). **d**, Isoform proportions (left) and total detected transcripts (right) per cell type reveal a clear switch in isoform usage: neuronal cell types express App695, whereas vascular and choroid plexus cells express App751 and App770. **e**, Spatial detection of *App* in a representative sample (P21F). **f**, *Clta* gene locus showing four isoforms defined by inclusion or exclusion of exons 5 and 6. **g**, *Clta* usage across samples shows an increase in mean detected transcripts per cell (middle) and a developmental switch in isoform usage from exon 5 inclusion to combined exon 5 and 6 inclusion (top and bottom). **h**, Mean isoform-level *Clta* expression per cell type shows a switch in neuronal populations from Clta_ex5 to Clta_ex5-6. Cajal-Retzius and medium spiny neurons continuously express Clta_ex5-6 from P1 to P21, whereas microglia and macrophages express isoforms lacking exons 5 and 6. **i**, *Clta* isoform usage per cell type is shown as proportions (left) and total detected transcripts (right). **j**, Spatial co-expression plots across development (P1-P21) comparing Clta_ex5 (red) and Clta_ex5-6 (green). **k**, Pseudobulk analysis of paired neuronal cell types (all neuronal cell types except MSNs and Cajal-Retzius cells) showing a major isoform switch from Clta_ex5 to Clta_ex5-6 between early postnatal time points (P1, P3 and P5) and later stages (P7, P14 and P21). The -log10(adjusted P value) is shown as a blue bar on the right side.

### Resolving isoform diversity with imaging-based transcriptomics (iSiT)

The β-amyloid precursor protein (*App*), whose proteolytic cleavage produces the Aβ peptide that accumulates in Alzheimer’s disease senile plaques^30^, also plays essential roles in normal neurodevelopment^31^. Although the *App* gene is already targeted in the Prime 5K panel, we designed isoform-specific probes targeting exon-exon junctions to resolve inclusion or skipping of exon 7, encoding the Kunitz-type protease inhibitor (KPI) domain, and exon 8, encoding the Ox-2 antigen domain, both conserved in human *APP*. Our design discriminates the three major *App* isoforms: App695 (lacking both domains), App751 (KPI only), and App770 (KPI+Ox-2) (**Fig.3a**). Brain-wide *App* transcript counts per cell increased over time, from an average of 2.8 molecules (P1) to 5.4 molecules per cell (P21), although overall isoform usage remained stable across the developmental time course (**Fig.3b**). At the cell-type level, isoform-specific analysis showed that App695 was predominantly expressed in neuronal populations, whereas the KPI-containing isoforms App751 and App770 were enriched in vascular and perivascular cell types^32^ (pericytes; vascular endothelial cells, VEC; vascular smooth muscle cells, VSMC; and vascular/leptomeningeal cells, VLMC) and in choroid epithelial cells (**Fig.3c-d**). Spatial mapping recapitulates this expression pattern, highlighting enriched App751/App770 signal in vascular endothelial cells as well as within the third and lateral ventricles, where choroid cells are located (**Fig.3e**). In the human brain, KPI-containing isoforms are elevated in Alzheimer’s disease and are thought to promote senile plaque formation^33^. In addition, cerebral hypervascularization separately contributes to cognitive decline and blood-brain barrier disruption during aging^34^. We therefore propose that mapping *App* isoforms in the aging brain would provide important insight into the etiology of Alzheimer’s disease.

We next examined clathrin, the principal structural component of clathrin-coated vesicles. The clathrin light chain gene *Clta* (CLCa/LCA) undergoes tissue- and developmental stage-specific alternative splicing of exons 5 (18aa) and 6 (12aa)^35^ generating isoforms that include either exon, both, or neither (**Fig.3f**). However, the cell type- and regional-specificity of these isoforms across the developing brain remains poorly understood. At the whole-brain level, *Clta* expression increased during development, accompanied by a switch from the exon 5-only to the exon 5-6 isoform, suggesting differential dynamic requirements for membrane trafficking (**Fig.3g**). Cell-type analysis provided additional resolution showing *Clta* expression predominates in neuronal populations, with the most pronounced isoform switch occurring between post-natal day P5 and P7. In addition, Cajal-Retzius neurons and medium spiny inhibitory GABAergic neurons (MSNs) of the caudate-putamen (CP) predominantly expressed the exon 5-6 isoform throughout development, whereas microglia and CNS macrophages predominantly expressed the isoform lacking both exons (**Fig.3h-j**). To further investigate developmental splicing regulation, we performed pseudobulk analysis, paired by neuronal cell types (excluding MSN and Cajal-Retzius) between early (P1, P3 and P5) and late (P7, P14 and P21) stages. This analysis identified 16 of 19 Prime 5K neuronal splicing regulators (**Fig.1i**) as significantly differentially expressed, consistent with the complex interplay of multiple factors that may contribute to developmental splicing regulation. Among these factors, *Elavl3* (HuC) and *Rbfox2*^36^ showed the strongest concordance, suggesting their contribution to *Clta* splicing dynamics during development (**Fig.3k**, **Supplementary Table 4** and **Methods**).

Together, our findings show that combinatorial usage of exon 5 and 6 tunes clathrin-mediated trafficking to developmental demands: the exon 5-only isoform supports membrane remodelling for axon guidance and neurite growth, the exon 5-6 isoform optimizes synaptic vesicle recycling and transmission^37^, wheras the isoform lacking both exons is restricted to microglia and CNS macrophages. Early expression of the exon 5-6 isoform in Cajal-Retzius and MSNs may therefore facilitate rapid circuit formation and network integration by promoting early synaptic stabilization and efficient incorporation into emerging neural networks, consistent with their established roles in early cortical circuit organization and striatal circuit integration^38^.

We next analyzed AMPA receptor subunit genes, glutamate-gated ion channels that mediate fast excitatory neurotransmission and are central to synaptic plasticity, circuit maturation, and cognitive function^39^. Dysregulation of AMPAR function and signaling is implicated in multiple neurological and neuropsychiatric conditions, including major depressive disorder^40^, Alzheimer’s disease^41^, epilepsy, and autism spectrum disorders^42^. Each AMPAR subunit undergoes alternative splicing to generate the flip and flop isoforms, which confer distinct channel kinetics, desensitization rates, and recovery from desensitization^43^. To resolve isoform usage in situ, we designed probes targeting both variants of each of the four AMPAR subunits (**Fig.4a**). Isoform-resolved profiling revealed marked cell-type- and stage-specific differences, with a progressive reduction in flip relative to flop isoform usage accompanying neuronal maturation (**Fig.4b**-**c**). Closer inspection of *Gria1* and *Gria2* transcripts uncovered pronounced regional heterogeneity in flip/flop expression across hippocampal and cortical regions. Notably, the flip isoform predominated in the superficial cortical layer (L2), whereas deeper layers (L3-6) primarily expressed the flop isoform (**Fig.4d**). Sub-clustering of excitatory neurons within Ammon’s horn further refined this pattern showing a striking subfield-specific switch: CA1 and DG neurons preferentially expressed the flop isoforms, whereas CA2 and CA3 neurons retained higher levels of the flip variants of *Gria1* and *Gria2* (**Fig.4e**).

We next sought regulators of flip/flop isoform expression in the hippocampus. Using the Zeisel et al. (2018) ClusterName annotation level, we observed a clear segregation of hippocampal pyramidal populations, with CA1 corresponding to TEGLU24 and CA3 to TEGLU23 (**Fig.4f**). Pseudobulk differential expression analysis of the matched whole-transcriptome single-cell reference identified *Celf4* and *Elavl2* (HuB) as enriched in TEGLU23, with *Elavl2* largely absent from TEGLU24 (**Fig.4g** and **Supplementary Table 5**). *Elavl2* is a neuron-specific RNA-binding protein that regulates alternative splicing during neuronal development and maturation and has been implicated in autism spectrum disorder (ASD) and related neurodevelopmental disorders^44^. Cell-type-specific expression and spatial density mapping at P21 confirmed this pattern, showing that *Elavl2* expression was largely mutually exclusive with the flop isoform and co-expressed with the flip isoform (**Fig.4h-i**). These findings reveal fine-grained spatial patterning of AMPAR isoform selection across distinct hippocampal circuits, suggesting that cell type-specific flip/flop splicing tunes synaptic kinetics and plasticity during postnatal maturation.

We uncovered additional cell-type-, developmental-stage- and region-specific patterns of isoform usage (**Extended Data Fig. 4**). This includes differential selection of exon8 and 8A in *Cacna1c*, inclusion of A1 and A2 exons in *Nlgn1*, skipping of exon3 in *Clstn1*, mutually exclusive usage of exons10a and 10b in *Dnm1*, regulation of *Nrxn1* SS4 domain, and differential usage of Z8 and Z11 exons in *Agrn*.

**Fig. 4:**
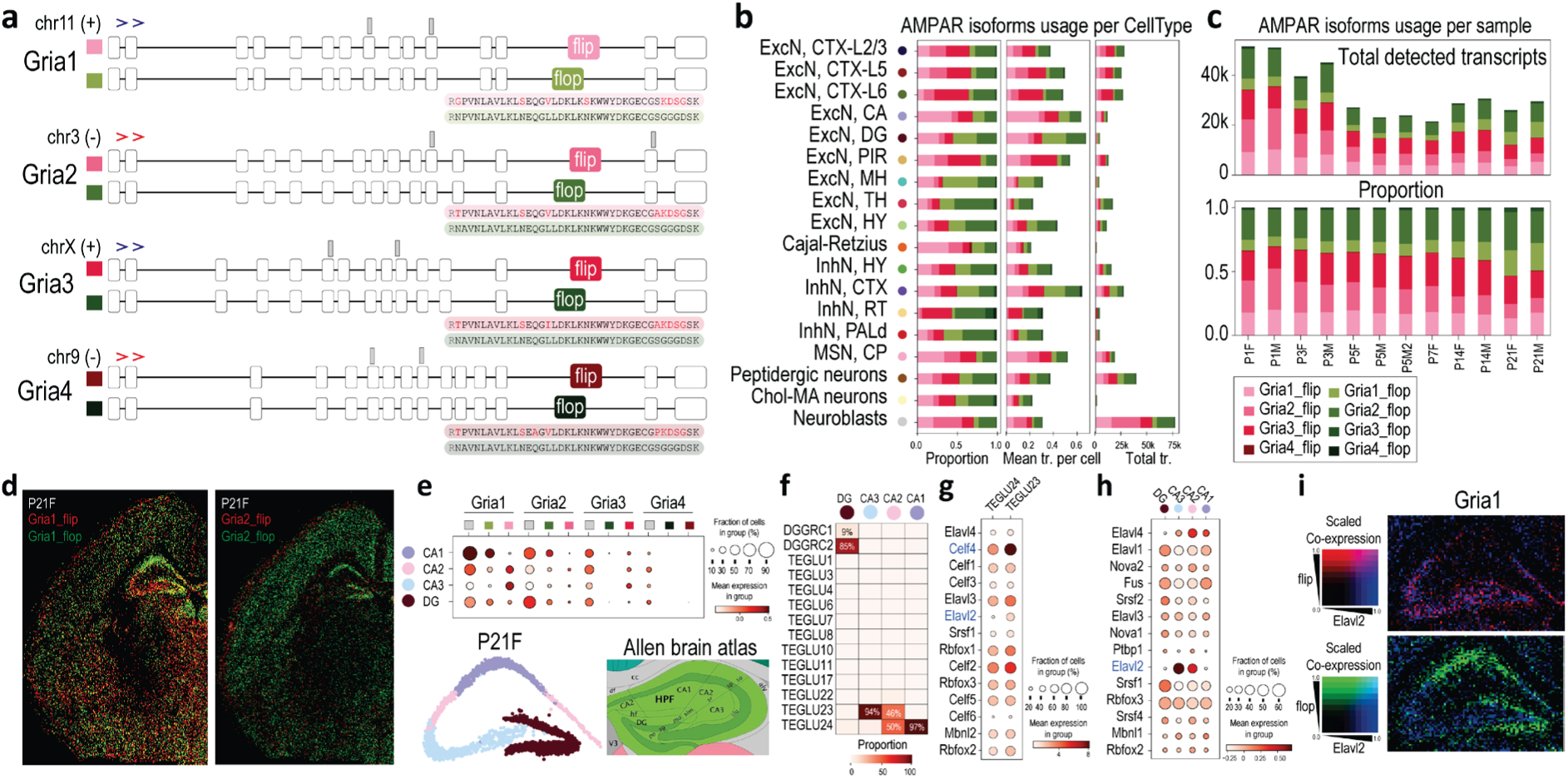
AMPA receptor isoform usage across cell type, space and time. **a**, Genomic loci of AMPA receptor subunits (*Gria1-4*), with gene-level Prime 5K probes indicated as grey rectangles. **b**, Flip/flop isoform expression across neuronal cell types, shown as proportional usage (left), mean number of detected transcripts per cell (middle), and total number of detected transcripts (right). **c**, Temporal dynamics of flip/flop isoform expression across development, shown as total detected transcripts (top) and proportional usage within each sample (bottom). **d**, Spatial co-expression (P21F) of flip and flop isoforms for *Gria1* (left) and *Gria2* (right). **e**, Sub-clustering of P21F excitatory neurons in CA and DG regions resolved CA1, CA2 and CA3 subfields and revealed distinct isoform expression patterns: the *Gria1* flip was enriched in CA2 and CA3, whereas the *Gria1* and *Gria2* flop isoforms predominated in CA1 and DG. **f**, Proportions of DG, CA1, CA2 and CA3 cells annotated as ClusterName from Zeisel et al. (2018). **g**, Expression of established master regulators of alternative splicing in TEGLU23 and TEGLU24 in the Zeisel et al., 2018, single-cell reference. **h**, Expression of established master regulators of alternative splicing in the Xenium dataset. **i**, Spatial co-expression of gene-level *Elavl2* and the flip and flop isoforms of *Gria1*.

**Fig. 5:**
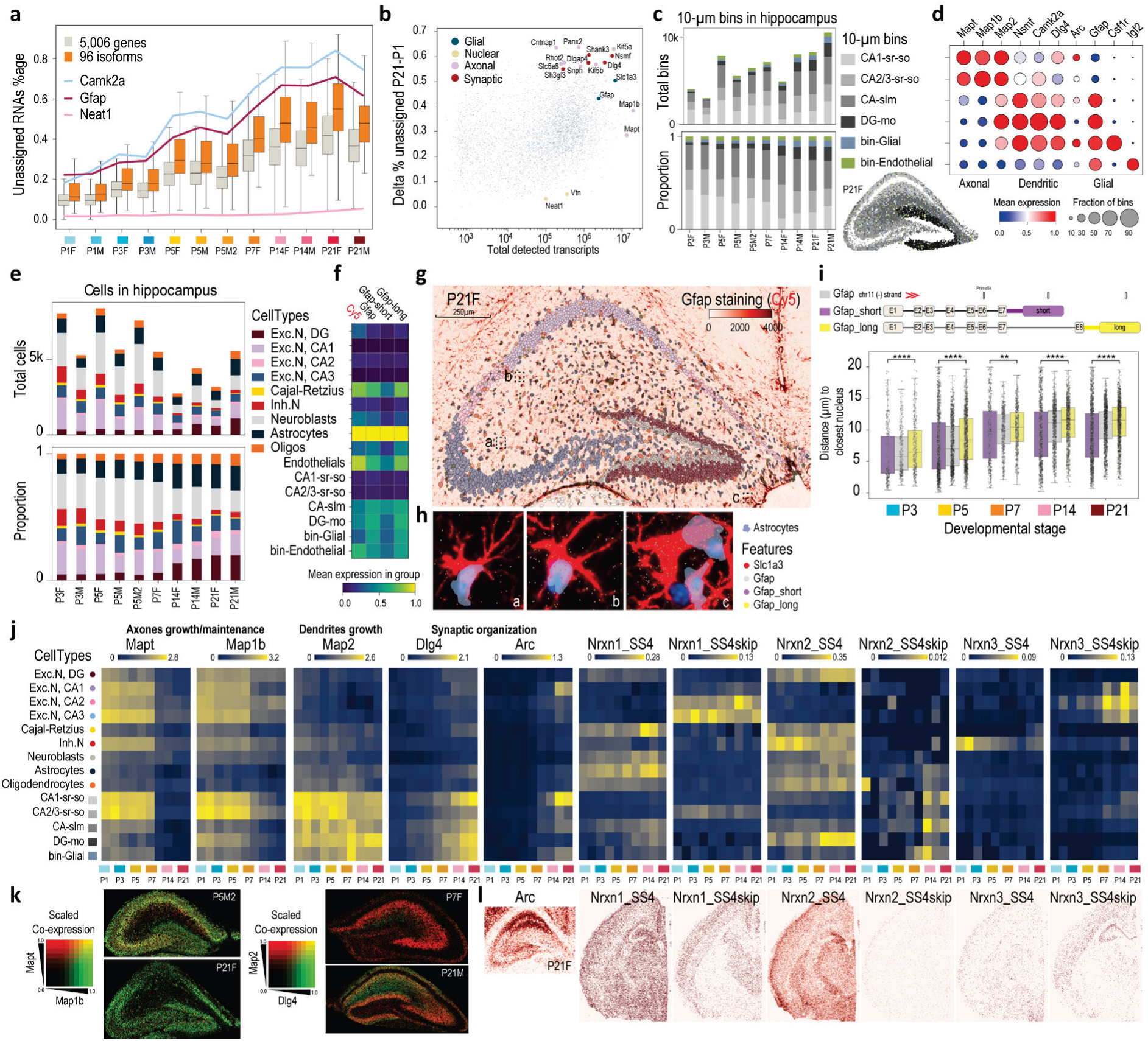
Unassigned RNAs (uRNAs) reveal isoform-level subcellular compartments. **a**, Box plots showing the percentage of unassigned RNAs per sample for the Prime 5K gene panel and the 96 isoform-level probes. Curves for selected genes are highlighted, including the nuclear transcript *Neat1*, the astrocyte marker *Gfap*, and the dendritically enriched gene *Camk2a*. **b**, Change (Δ) in the percentage of unassigned RNAs between P21 and P1 stages as a function of the total number of detected transcripts across samples. Representative genes associated with glial, nuclear, axonal and dendritic/synaptic compartments are indicated. **c**, Total number (top) and proportion (bottom) of 10-μm spatial bins in the hippocampal region per sample, colored by bin type. Bottom right, spatial map of bin types in the P21F sample. **d**, Dot plot showing expression of representative marker genes across bin types. **e**, Total number (top) and proportion (bottom) of cell types identified in the hippocampal region. **f**, Mean expression of *Gfap* measured by protein staining (Cy5), gene-level transcripts and isoform-level transcripts (short and long isoforms). **g**, Spatial distribution of *Gfap* protein staining and cell types in the hippocampal region of the P21F sample. **h**, Zoom-in views of three regions from (g), showing astrocyte segmentation based on Xenium multimodal data and *Gfap* staining delineating real astrocyte boundaries. Detected transcripts for *Slc1a3*, *Gfap* (gene-level) and *Gfap* isoforms are overlaid. **i**, Top: schematic of the *Gfap* locus indicating gene-level and isoform-level probesets. Bottom: box plot showing the distribution of distances between detected *Gfap* transcripts and the nearest astrocyte nucleus centroid, highlighting that short-isoform transcripts are located closer to nuclei than long-isoform transcripts. **j**, Left: mean expression per cell type and bin type across developmental time points for representative axonal (*Mapt*, *Map1b*), dendritic (*Map2*) and synaptic organization (*Dlg4*, *Arc*) genes. Right: expression levels of SS4+ and SS4− isoforms of neurexins (*Nrxn1*, *Nrxn2*, *Nrxn3*). **k**, Spatial co-expression plots at 5µm binning (green-to-red colormap) corresponding to representative events shown in (j). **l**, Spatial visualization (P21F) of expression levels (red colormap) corresponding to representative events shown in (j).

**Fig. 6:**
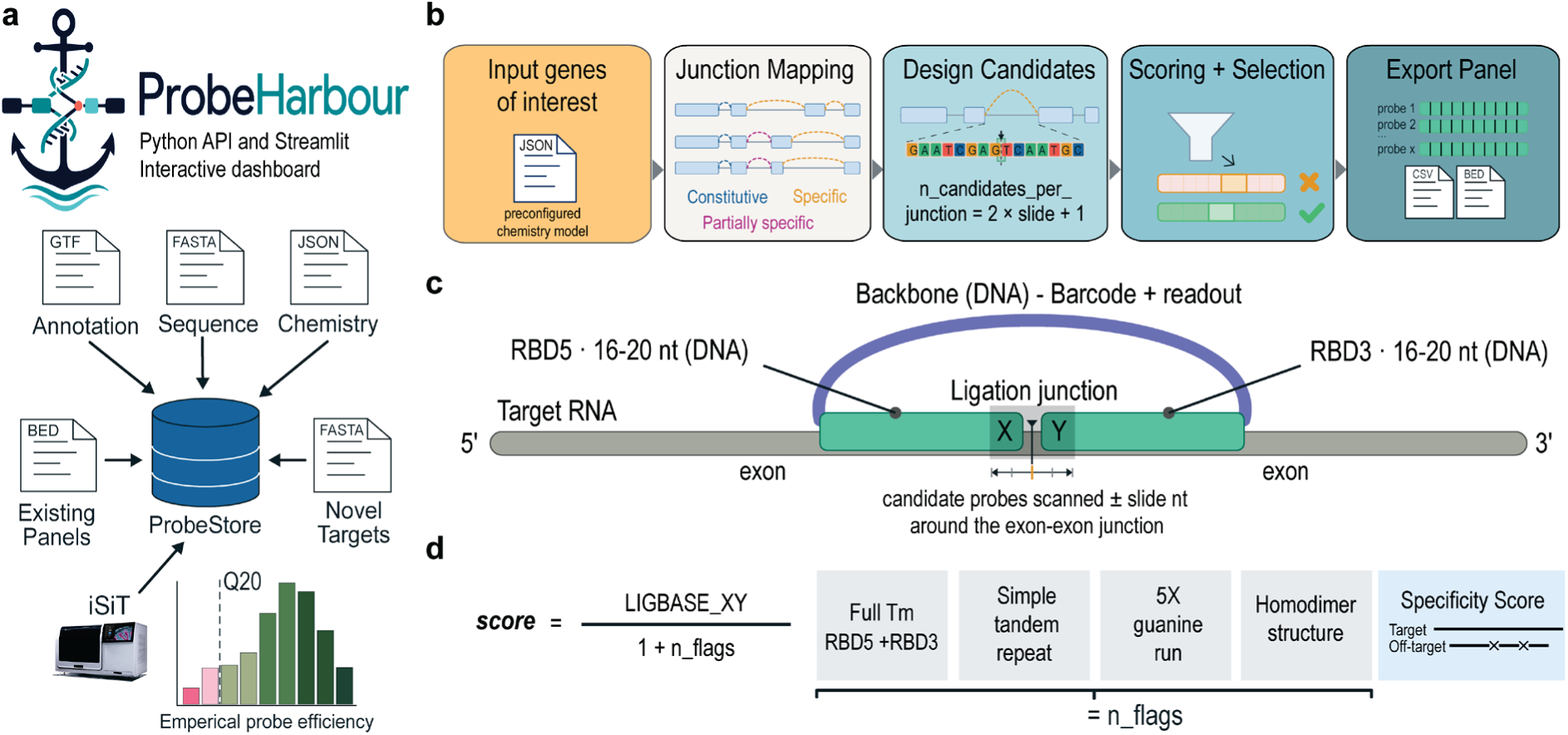
ProbeHarbour enables isoform-resolved spatial transcriptomics probe designa. **a**, ProbeHarbour integrated Python API, organised around the ProbeStore data structure. Inputs are a genome annotation (GTF), reference sequence (FASTA) and a chemistry model (JSON), with optional existing panels (BED) and novel targets (FASTA). Empirical per-probe efficiency (QV) from iSiT experiments is fed back into the ProbeStore to inform future designs. **b**, Schematic of the ProbeHarbour design workflow, driven through the Python API and equally accessible from the interactive Streamlit dashboard: input genes are mapped to constitutive, specific and partially specific junctions, candidate probes are generated across a sliding window (n_candidates_per_junction = 2 x slide + 1), scored and selected, and exported as CSV or BED panels. **c**, Representative imaging-based visualization of padlock probe architecture: two DNA arms, RBD5 and RBD3 (16-20 nt), flank the exon-exon ligation junction on the target RNA and carry the barcode/readout backbone; candidate probes are scanned +/- slide nt around the junction. **d**, Candidate probe scoring strategy used to optimize chemical efficiency and overall performance. The raw score is LIGBASE_XY / (1 + n_flags); the annotation flags that modulate the ligation-dinucleotide preference weighting (full-binder Tm, simple tandem repeat, five-guanine run, homodimer structure) are shown on the right, together with the specificity score. The scheme is fully configurable through the JSON, so a user-defined score can be built from the per-probe features in place of this default.

### Resolving isoform diversity at subcellular resolution

Across all samples, we detected ∼2.2 billion high-quality transcripts (QV≥20). Of these, ∼1.5 billion (69.2%) were assigned to cell segmentation masks, whereas ∼0.7 billion (30.8%) remained unassigned (uRNAs) (**Fig.1d**). Notably, the fraction of uRNAs progressively increased from 11.3% at P1 to 49.1% at P21, with marked gene-specific differences. This trend likely reflects the increasing morphological complexity of differentiated cell types, including neurons, astrocytes, and oligodendrocytes, which develop specialized cellular processes extending beyond conventional soma-based segmentation boundaries. For example, the predominantly nuclear transcript *Neat1* showed consistently low uRNA fractions across developmental stages. In contrast, *Camk2a*, enriched in dendrites, and the astrocyte marker *Gfap* exhibited substantially higher uRNA fractions, which increased with brain maturation (**Fig.5a**). Comparative analysis between P1 and P21 showed increased uRNA fractions for synaptic genes, including *Dlg4* (PSD-95), *Shank3*, *Dlgap4* and *Sh3gl3*, and the axonal motor proteins *Kif5a* and *Kif5b*, consistent with synaptic localization. Glial transcripts, including *Gfap* and *Slc1a3*, also showed increased unassigned fractions, whereas nuclear (*Neat1*) and perinuclear (*Vtn*) transcripts remained largely assigned to cells throughout brain maturation (**Fig.5b**).

We therefore hypothesized that a substantial portion of these uRNAs reflect specific subcellular RNA organization, potentially linked to region-specific synaptogenesis. To investigate this, we aggregated uRNA signals into 10-µm bins across hemispheres at P3, P5, P7, P14 and P21. We observed a progressive increase in both bin number per sample and transcripts per bin (**Extended Data Fig. 5a-c**). Unsupervised clustering of bins revealed two classes of regionally organized community: those with cell-type-specific expression signatures, consistent with incomplete segmentation of nearby cells, and those with distinct profiles, consistent with distal subcellular compartments. Gene set enrichment analysis identified clusters associated with neuronal projections and synaptic components (**Extended Data Fig. 5d-f**). We focused on the hippocampus, which showed the highest uRNA density and is a canonical model for synaptic biology. Clustering of hippocampal uRNAs identified four spatially distinct neuropil compartments matching known subregions: CA1 stratum oriens/radiatum (CA1-sr-so), CA2/3 stratum oriens/radiatum (CA2/3-sr-so), stratum lacunosum-moleculare (CA-slm) and the dentate gyrus molecular layer (DG-mo) (**Fig.5c-d** and **Extended Data Fig. 5g**). We performed pseudobulk comparisons between pyramidal neuron (somata) and corresponding dendritic compartments (neuropil) in CA1, CA2/3 and DG (**Extended Data Fig. 5h-k**, **Supplementary Tables 6-9** and **Methods**). Incorporating cell-type annotations across hippocampal subregions revealed enrichment of Cajal-Retzius in the DG-mo, consistent with their role in hippocampal lamination via Reelin signaling. This enrichment declined postnatally. In parallel, inhibitory interneurons decreased in CA1-sr-so and CA2/3-sr-so, while neuroblasts progressively transitioned into DG granule neurons (**Fig.5e**).

We next focused on astrocytes, key regulators of synapse formation and maturation through the secretion of synaptogenic factors^45^. Post-Xenium *Gfap* protein staining showed high concordance with unassigned as well as cell-assigned *Gfap* transcripts, indicating that standard segmentation fails to capture astrocyte boundaries and their highly ramified protrusions (**Fig.5f,g**). At the isoform level, Gfap-long detected transcripts show a greater mean distance to the closest astrocyte centroid than Gfap-short transcripts, consistent with their enrichment in peripheral processes^46^ (**Fig.5h,i** and **Methods**)

To orient the individual hippocampal subcellular compartments, we mapped canonical markers of axon growth and maintenance (*Mapt*, *Map1b*) mainly expressed before P14 in sr-so compartments, dendritic development (*Map2*), and synaptic organization (*Dlg4*, *Arc*) increasing over the developmental time course. At the isoform level, Neurexins, known as presynaptic adhesion molecules mediating trans-synaptic coupling with Neuroligins, showed complex alternative splicing. SS4 splicing modulates ligand binding and thereby presynaptic release-site composition and alignment with postsynaptic NMDA- and AMPA-receptor-enriched dendritic scaffolds^47^. SS4-skipped *Nrxn1* (all timepoints) and *Nrxn3* (P14-P21) were enriched in CA1/2/3 pyramidal neurons, whereas SS4-including isoforms showed broader expression in DG granule neurons, Cajal-Retzius cells, neuroblasts (*Nrxn1*/*Nrxn2*) and early interneurons (*Nrxn3*). Consistently, SS4-including isoforms were enriched in synaptic layers (CA-slm, DG-mo), where entorhinal axons form maturing dendritic synapses during late postnatal development (P14-P21). Collectively, these findings support that Neurexins isoforms contribute to compartment-specific presynaptic programs associated with the progressive specialization of axon-dendrite connectivity during hippocampal maturation (**Fig.5j-l**).

### ProbeHarbour: an open framework for isoform-resolved probe design

Although our initial isoform-level add-on panel design provided high spatial and temporal resolution for the majority of genes examined, 27 out of 96 probes exhibited a median QV below 20, precluding the use of the full panel (**Extended Data Fig. 6a**). This highlights the need for methodological improvements and probe design optimization, particularly for single-probe-per-probeset strategies targeting exon-exon junctions, which impose design and efficiency constraints. To address these limitations, we developed ProbeHarbour, a Python package enabling the systematic design and selection of isoform-resolved probes. Starting from a preconfigured Xenium chemistry model JSON (v1, Prime 5K), users define a set of genes of interest. Based on the GENCODE gene model, ProbeHarbour iteratively evaluates each gene, identifying constitutive, partially specific and specific junctions. For each user-selected junction of interest, ProbeHarbour scans candidate probes across a sliding window around the exon-exon boundary and computes a series of metrics capturing the key parameters governing probe hybridization efficiency and specificity (**Methods**) for candidate scoring and optimal selection. At the end of the workflow, users can review and export CSV and BED files that are fully compatible with the 10x Genomics cloud environment for validation and synthesis. ProbeHarbour provides both an API and an interactive dashboard (**Fig.6a-c**).

To establish a composite probe-design score, we reverse-engineered publicly available Xenium predesigned panels (Prime 5K and V1, mouse and human; **Extended Data Fig. 6b**) to identify the sequence and chemistry features associated with optimal probe performance. From the predesigned probes we derived the thermal design windows from full-binder and per-arm melting temperature distributions ([71,77]°C for Prime 5K, [68,82]°C for V1; **Extended Data Fig. 6c**), and the ligation-dinucleotide preference weights from junction usage under active design rules, separately for each chemistry (**Extended Data Fig. 6d**). These features were combined with secondary-structure, repeat-content and probe-to-transcript specificity terms into a composite raw score, LIGBASE_XY / (1 + n_flags), where LIGBASE_XY is the weight of the ligation dinucleotide at the cut site and n_flags counts the sequence and thermodynamic penalties incurred by the probe (**Fig.6d**). The scoring scheme is not fixed: the flags, the ligation-dinucleotide weights and the score expression itself are all defined in the JSON configuration, so beyond this default composite score users can specify a custom, feature-based score built from any of the per-probe metrics.

The resulting ProbeHarbour score reproduced the vendor design rank almost exactly (n=95, Spearman rho=0.995; **Extended Data Fig. 6e**), confirming that the reverse-engineered model recovers the proprietary scoring at the level of the score itself, and not only in which probes are selected. The score correlated only weakly, however, with the median QV observed in our iSiT experiments (n=96, Spearman rho=0.242; AUROC=0.680 for QV≥20; **Extended Data Fig. 6f**). Design-time sequence and thermodynamic features therefore act as an effective quality filter but do not by themselves predict in situ probe efficiency, indicating that additional determinants govern probe performance. ProbeHarbour was designed to close this gap. The median QV measured for every probe in an iSiT run is recorded back into the ProbeStore, so that empirical probe efficiencies accumulate across experiments and can be used to re-weight the design parameters. This feedback loop will support data-driven refinement of junction-probe design as isoform-targeted panels scale on emerging platforms such as ATERA.

Together, these analyses establish ProbeHarbour as both a characterisation framework for existing Xenium probe panels and a design framework for isoform-aware spatial transcriptomics.

## Discussion

In this study, we established imaging-based spatial isoform transcriptomics (iSiT), a framework that extends imaging-based spatial transcriptomics beyond gene-level measurements by enabling isoform-resolved profiling on the Xenium platform. By integrating large-scale gene expression measurements with targeted isoform detection, iSiT enables spatial characterization of alternative splicing events at cellular and subcellular resolution requiring a single probe per target enabling analysis of internal transcript structure. Applied to the developing mouse brain, iSiT revealed extensive spatial and temporal regulation of transcript isoforms, encompassing developmental isoform switching, exemplified by *Clta* in neurons; cell-type-specific isoform usage, including *App* in endothelial and choroid cells or AMPAR flip/flop isoforms in distinct hippocampal cell populations. Beyond differential isoform usage, iSiT captured subcellular localization patterns of alternatively spliced transcripts, including neurexins in hippocampal compartments and *Gfap* isoforms in astrocytes. These results demonstrate that spatial isoform profiling uncovers regulatory layers that remain inaccessible to conventional gene-centric approaches. To facilitate broader adoption, we developed ProbeHarbour, a computational toolkit for designing isoform-specific probes compatible with Xenium chemistry. Together, these experimental and computational advances establish iSiT as a scalable approach for isoform-resolved spatial transcriptomics, opening new avenues to investigate how alternative splicing shapes cell identity, subcellular organization and developmental programs.

Several limitations of the current implementation must be considered. Because iSiT detects predefined exon-exon junctions, it resolves specific alternative splicing events rather than full-length transcript architectures, as achieved with long-read sequencing-based approaches. Accordingly, cassette exon inclusion or exclusion and alternative 5′ or 3′ UTR usage cannot always be unambiguously assigned to a single full-length transcript identity. Although previously unannotated isoforms may represent an additional source of ambiguity, their presence is expected to be limited in healthy tissues^9^. We therefore interpret our findings conservatively, focusing on specific splicing events and their predicted effects on protein composition, localization, and regulatory properties.

Although the present implementation was demonstrated using a limited targeted panel of 96 isoforms, the framework is inherently scalable. Future imaging-based chemistries, such as ATERA, are expected to combine genome-scale gene expression profiling (∼18,000 genes) with expanded isoform-targeted panels (∼1,000 custom targets) extending spatial alternative splicing analyses from targeted biological programs toward comprehensive regulatory landscapes. iSiT therefore offers a practical route toward isoform-resolved spatial transcriptomics. By overcoming the biases of poly(A)-based single-cell and in situ capture approaches, iSiT will improve access to long transcript isoforms that remain underrepresented in conventional reference datasets, thereby providing a richer molecular foundation for future virtual cell models^48^.

While demonstrated here in the developing mouse brain, iSiT is readily applicable to diverse tissues and biological systems compatible with Xenium imaging. In the nervous system, isoform regulation plays a central role in development and disease^49^, and antisense oligonucleotide strategies have highlighted the therapeutic potential of restoring physiological isoform states^50^. By enabling spatial mapping of transcript diversity, iSiT provides a scalable framework to determine how isoform dysregulation contributes to neurodevelopmental and neurological disorders. We anticipate that imaging-based spatial isoform profiling will become an essential dimension of next-generation spatial transcriptomics, enabling routine investigation of transcript diversity at cellular and subcellular resolution.

## Online Methods

### Mice breeding

Four female and two male C57BL/6N mice obtained from Charles River were used as breeding animals. Their offspring were used for the experiments described below. In total, 12 pups were analysed: one male and one female at each of six postnatal stages (P1, P3, P5, P7, P14 and P21). All animal care and experimental protocols were conducted according to European, national and institutional regulations (Protocol numbers: 00236.03, IPMC approval F0615252). Personnel from the laboratory performed all experimental protocols under strict guidelines to ensure careful and consistent handling of the mice. The animals were maintained under a 12-h light-dark cycle with free access to food and water.

### Brain collection and tissue preparation

C57BL/6N Mice postnatal day P1, P3, P5, P7, P14 and P21 were sacrificed by cervical dislocation. After extraction, brains were immediately embedded in O.C.T. Embedding Matrix for Frozen Sections (KMA010000A, Leica) and stored at -80°C until use. Tissue processing followed the manufacturer’s protocol (10X Genomics, CG000579 Xenium In Situ - Fresh Frozen Tissue Preparation Handbook, Rev F). Using a cryo-microtome (Leica CM3050), 10-µm-thick hemispheric coronal brain sections (Bregma ∼-1.94 mm) were collected on Xenium Spatial Gene Expression slides (3000941, 10x Genomics). For each developmental stage, two sections from two different mice were used; the two replicates were collected on two different Xenium Spatial Gene Expression slides and processed in parallel.

### Isoforms 100 add-on panel design

Complementing the predesigned 5,006-gene Xenium Prime 5K Mouse Pan Tissue & Pathways Panel we designed a 100 add-on panel, of which 4 code words were used for gene-level splicing factors (*Nova1*, *Rbm4*, *Xbp1* and *Mbnl3*) and 96 code words to target specific isoform splice junctions for 45 synaptogenesis key player genes. For each junction of interest we design all potential probes (i.e. 16 nt rbd3 + 16 nt rbd5) spanning the junction authorizing a maximum of 6 nt offset before or after the junction. For optimal probe selection we evaluate the following criteria: (i) rbd3/5 Tm [50-70°C], and combined full Tm [70-82°C]; (ii) avoid repeat and poly-N (i.e. > 5); (iii) prefer optimal [AT-TA-GA-AG] and neutral [AA-TC-CA-TT-TG], avoid unfavorable/non-preferred dinucleotide ligation junction; (iv) avoid off-target matches in whole mouse transcriptome using blast; (v) avoid probe closer than 75 nt from a Xenium gene-level probe. To prevent optical crowding issue, we excluded seven highly expressed genes (*Myl6*, *Ctsd*, *Mbp*, *Calm2*, *Slc1a2*, *Plp1* and *Aldoa*) from the panel design, based on high gene-level tp10k per cell type across three reference mouse brain single-cell atlases. Out of our 45 final gene targets, 17 were absent and 28 were already profiled by Xenium Prime 5K at the gene-level with one (4), two (20) or three (4) different probesets. We therefore expect a reduced sensitivity at the isoform-level for those 28 genes due to RNA molecules sequestered by the gene-level probesets. For all 45 genes, we target auto-exclusive junctions to avoid interrogating the same mRNA molecule with two concurrent Isoform-level probes. Therefore, we consider our isoform-level add-on panel as a boost of one additional single-probe probeset for the 28 genes and the adding of new unique single-probe probeset detection for the 17 other genes. For those 28 genes, we defined a code word avoiding interrogating the same 5 cycles as the gene-level probesets. During probe design, the flip and flop annotations of the AMPA receptor probes were inverted. This was corrected in all files and objects provided as raw and analyzed data using the Xenium Ranger relabel pipeline with an updated JSON.

### 10x Genomics Xenium

We used the predesigned Xenium Prime 5K Mouse Pan Tissue & Pathways Panel (PN-1000725) complemented by our Custom 100 gene Add-On isoform-level Panel (Design ID 3J6VCV). Twelve mouse brain sections were profiled by the IPMC genomic functional platform UCAGenomiX, member of the French Genomics National infrastructure. Probe hybridization, ligation and rolling circle amplification, with multimodal cell segmentation and staining, were performed following the manufacturer’s protocol (CG000581 RevE and CG000760 RevC, 10x Genomics). Background fluorescence was chemically quenched. Imaging and signal decoding were done using the Xenium Analyzer instrument following manufacturer’s user guide (CG000584 Rev K, 10x Genomics).

### Gfap protein staining

For post-staining 10x Genomics Xenium slides, no additional fixation or permeabilization was required. Slides were incubated with Animal-free blocker buffer (SP-5030-250, Vector laboratories) for 1h at room temperature. Primary antibodies (anti-Gfap, Invitrogen, #13-0300) diluted in the blocking buffer at 1/200, was applied and incubated overnight at 4°C. The slides were then washed three times with PBS-T before adding the secondary antibodies (anti-rat-Atto647), diluted 1/500 in the blocking buffer and incubated for 1 h at room temperature. Following another set of three washes with PBS-T, the slides were mounted with FluoroMount-G with DAPI solution (Invitrogen), and stored at 4°C until imaging. Acquisition was performed using an Axioscan7 (Zeiss) with a 20x air objective.

### 10x Genomics Xenium data processing

The 12 sections included in this study were formatted as SpatialData^51^ objects (v0.7.2) and analyzed as AnnData objects using Scanpy^52^ (v1.12) and Sparty (https://github.com/cobioda/sparty). Cell-by-gene matrices from Xenium Ranger were normalized and log-transformed. Neighborhood graphs were computed using 40 principal components and 15 nearest neighbors. Datasets were integrated with Harmony (rapids-singlecell^53^), followed by Leiden clustering. Cell types were annotated by label transfer using scMusketeers^54^ and an adolescent mouse central nervous system reference^4^ at both TaxonomyRank4 (T4) and ClusterName (CN) levels. Automated label transfer was performed five times independently at each annotation level, and majority voting was used to assign T4 and CN labels to each cell. Label transfer robustness across the five runs was recorded in the final Anndata object as T4n for TaxonomyRank4 and CNn for ClusterName denoting the number of times one T4 or CN were assigned to each cell. Final cell type (CellType) annotations were assigned based on Harmony-integrated Leiden clusters, T4 and CN labels, and spatial context. For cell type-specific expression heatmaps, we further excluded putative doublet cells arising from inaccurate cell segmentation identified using Scrublet^55^ (doublet_score cutoff > 0.2), excluding 41,349 cells. We further excluded cells showing a non-robust automated T4 assignment (T4n in [1,2]; 65,194 cells), a cluster of spatially unresolved excitatory neurons (6,997 cells), and a cluster of hypendymal cells (22 cells). The resulting filtered AnnData object, containing 1,234,789 cells, was used for all downstream analyses and data visualizations presented in the manuscript.

### *Gfap* protein staining alignment and quantification

Acquired *Gfap*-stained OME-TIFF image files were aligned to Xenium morphology images using Xenium Explorer. The resulting alignment matrices were exported and subsequently applied to SpatialData objects for each individual sample using the Sopa^56^ “explorer add-aligned” pipeline (v2.2.2). Per-cell quantification was performed with the Sopa aggregate_channels function, using the cell_boundaries segmentation mask and the aggregation mode set to "average".

### Cell-type proportion analysis

Each sample was rotated to match a standard left-right coronal section orientation. A single representative hemisphere was then extracted per animal by visual inspection, using x-axis boundaries defined by alignment of the third ventricle regions. The replication unit for all compositional statistics is therefore the individual mouse: each animal contributes exactly one hemisphere, and left and right hemispheres are never treated as separate replicates. We identified a spatial mismatch in the P1M sample relative to the other mid-hippocampal sections (-1.6 to -2.5 mm from bregma), with P1M falling outside this range according to the developmental mouse brain CCF^57^; we therefore excluded it from cell-type proportion analyses. The remaining 11 animals were grouped into three developmental windows and compared using independent two-sample t-tests, with each animal pooled across sex on the basis of the male/female concordance shown in Ext.Data.Fig.2c,d: P1-P3 (n=3 animals: P1F, P3F, P3M), P5-P7 (n=4 animals: P5F, P5M, P5M2, P7F) and P14-P21 (n=4 animals: P14F, P14M, P21F, P21M).

### Pseudobulk differential expression analysis

Differential gene expression analysis presented in the manuscript was performed using the PyDESeq2^58^ (v0.5.4) and decoupler^59^ (v2.1.6) pseudobulk strategy using standard default options (>100 counts, 5 cells, and 10 genes per aggregate) implemented within Sparty package tl.pseudobulk method. Multiple testing was corrected using the Benjamini-Hochberg false discovery rate (FDR). Differentially expressed genes were defined as those with an adjusted p-value (FDR) ≤ 0.05 and an absolute log2 fold-change ≥ 1. Figure 3-k compares early (P1, P3 and P5) versus late (P7, P14 and P21) stages, paired by neuronal cell types, excluding MSNs and Cajal-Retzius. Extended Data Figure 5 panels h, i, j, and k compare, respectively, CA1 pyramidal neurons (CA1-sp) and bin-CA1-sr-so; CA2/3 pyramidal neurons (CA2/3-sp) and bin-CA2/3-sr-so; DG granule neurons (DG-sp) and bin-DG-mo; and bin-CA1/2/3-sr-so between P3-P7 and P14-P21, all grouped by individual sample. For the bin-versus-cell comparison conducted within the same donor and the stage comparisons conducted within the same cell type, paired designs were used to account for inter-individual variability (∼donor + CellType) and (∼CellType + stage), respectively.

### Gene and Isoform -level expression heatmaps

All heatmaps were generated from filtered AnnData objects. Raw transcript counts per cell were normalized to the median number of detected transcripts per cell and log-transformed using Scanpy preprocessing functions (pp.normalize_total and pp.log1p). Heatmaps were then produced using Seaborn’s heatmap function with the cividis colormap. The vmax parameter was manually adjusted to optimize visualization of expression ranges across cell types and samples.

### Spatial density and co-expression plots

All detected transcripts, including both cell-assigned and unassigned RNA, were included in the spatial density and co-expression analyses presented in the manuscript. Spatial density plots were generated using the Sparty package (pl.plot_density) with a bin size of 20µm binning and a pct_max threshold of 0.98, such that the top 2% of values were capped to the maximum intensity. Spatial co-expression analyses were performed using Sparty package (pl.colocalization), using the same bin size and pct_max threshold. Expression values were scaled between 0 and 1 using the highest pct_max value observed across the two genes or isoforms compared (default, scale=common), or independently by feature (scale=independent). Signals were visualized using additive RGB color encoding, with red and green by default and blue used for splicing regulators. Co-localization within bins resulted in additive color mixing (e.g., yellow for overlapping red and green signals).

### Hippocampal region extraction

Hippocampal boundaries were defined using dentate gyrus (DG) and Ammon’s horn (CA) excitatory neurons. First, a spatial neighborhood graph was computed using Sopa spatial_neighbors (percentile = 95). Secondly, an alpha-shape geometry was generated from these cells and their neighborhood graph using the Sopa geometrize_niches function. Thirdly, the resulting geometry was refined by applying a convex hull operation followed by a 50μm buffer to ensure inclusion of all cells within the hippocampal region. Finally, the hippocampal polygon was used with the SpatialData polygon_query function to subset each spatial dataset to the hippocampal region of interest. Hippocampal region was subsequently used for quantification of unassigned RNA molecules at the spatial bin level, for each individual sample.

### Unassigned RNA bins definition

For each sample, unassigned RNA molecules were quantified using Sparty tl.unassigned_RNA function. This procedure generated a new layer of spatial shapes (bin_boundaries), a new layer of tables (bin_unassigned) and a corresponding count matrix (bin x features). Bins were generated at a resolution of 10μm, and only bins containing at least 100 detected transcripts were retained for downstream analyses. *Gfap* staining was quantified for each spatial bin using the Sopa aggregate_channels function, using the bin_boundaries and the aggregation mode set to "average". The “bin_unassigned” AnnData objects from all samples were merged into a single AnnData object and were processed using a standard single-cell analysis including normalization, PCA, Harmony batch correction, and Leiden clustering, using rapids-singlecell package. The integrated and annotated cell- and bin-level datasets were then combined for differential expression analyses and visualization. Cell and bin annotations were also transferred back to the corresponding SpatialData objects to enable integrated spatial visualization of both segmentation-derived cells and unassigned RNA bins.

### *Gfap* -long versus -short isoforms distance to nucleus analysis

For each sample, transcript-to-nucleus distances were computed using the hippocampal SpatialData objects. High-quality decoded transcripts (QV≥20) corresponding to *Gfap*, Gfap-long, and Gfap-short, including both Astrocyte-assigned and unassigned transcripts, were mapped to their nearest Astrocyte nucleus centroid, and transcript-to-nucleus distances were computed using Euclidean distance. A maximum distance threshold of 20μm was applied, and transcripts located beyond this distance from their nearest Astrocyte nucleus were excluded from subsequent analyses. For statistical analyses, only Astrocyte nuclei containing at least one Gfap-short transcript and at least one Gfap-long transcript were retained, thereby restricting the analysis to paired cells expressing both isoforms. For each retained cell, the median transcript-to-nucleus distance was calculated separately for the two isoforms. Distance comparisons were therefore performed on paired observations. For each developmental stage (P3, P5, P7, P14, and P21), paired median distances between the two isoforms were compared using two-sided paired Wilcoxon signed-rank tests. P-values were adjusted for multiple testing using the Benjamini-Hochberg false discovery rate (FDR) correction implemented with the multipletests function.

### 10x Genomics Visium HD

Ten-micrometer sections were cut using a Cryostat and placed directly onto a Superfrost Plus microscope slides. Slides were stored at -80 °C into slide mailers with desiccant until use. The Visium HD slides were prepared as described in the Visium HD 3’ Fresh Frozen Tissue Preparation Handbook (CG000804 Rev A, 10X Genomics) and followed by the 10x Genomics Visium HD 3’ Spatial Gene Expression User Guide (CG000805 Rev B, 10X Genomics) with CytAssist. Libraries were sequenced on Illumina NextSeq2000 according to 10x Genomics guidelines (paired-end reads 43b x 75b). For long-read sequencing, full-length cDNAs were size-selected using a 0.50x SPRI Select procedure, ONT adapters were then ligated, and the libraries were sequenced using one PromethION flow cell for 72 hours.

### 10x Genomics Visium HD data processing

Short-reads were processed using 10x Genomics Space Ranger workflow (4.0.1) using the GRCm39-2024-A reference. Long-reads were processed using epi2me-labs/wf-single-cell Nextflow workflow (3.3.4) using the Gencode_mouse_M38_GRCm39 reference after Percula pipeline. Quantification file were then loaded as SpatialData objects and analyzed at 8µm binning resolution

### ProbeHarbour API and dashboard

ProbeHarbour is a Python package for designing exon-exon junction probes that discriminate isoforms of the same gene under 10x Genomics Xenium chemistry. Parameters for design, scoring and selection are supplied as versioned JSON rulesets per chemistry and species, so that every candidate carries the identity of the configuration under which it was designed. The API is organised around ProbeStore, a serializable container holding one row per probe with sequences, coordinates, thermodynamics, score components, specificity, panel compatibility, selection status and provenance. Existing panels can be reconstructed from BED12 and reference FASTA, so that an add-on panel is evaluated alongside the panel it will accompany; stores can be subset, merged, rescored and exported as BED, CSV and FASTA. The same workflow is available through a Streamlit dashboard.

Junctions in the supplied annotation are classified as constitutive when present in all annotated transcripts, specific when present in exactly one, and partially specific otherwise; annotation depth makes this a conservative lower bound. Candidates are generated by moving the ligation cut across a window around each junction: for cut position c and arm length a, RBD5 = sequence[c−a:c], RBD3 = sequence[c:c+a], and the ligation dinucleotide is the last base of RBD5 followed by the first of RBD3 in transcript-sense orientation. A symmetric range of ±s yields up to 2s+1 candidates per junction, deduplicated across transcript models by arm sequences and junction identity, with arms mapped to strand-aware genomic coordinates so that BED12 export excludes the intron. Prime 5K workflows use 16 nt arms, V1 workflows 20 nt. Melting temperatures, GC content, hairpin, homodimer and heterodimer free energies are computed with Primer3 (v2.1) (80 mM monovalent, 5 mM divalent, 0 mM dNTP, 1,500 nM oligonucleotide, Schildkraut correction, 50 °C structure evaluation). Two sequence features are flagged: a 3-4 nt motif repeated at least three times in the complete binder, and a run of five or more cytosines in either arm. Specificity is assessed by BLAST against the species cDNA, run on the complete binder and on each arm, and candidates within 75 nt of an existing gene-level probeset are flagged.

### Exon-exon junction design and characterization

Published Xenium panels were reconstructed from 10x Genomics BED12 locations and matching reference FASTA: Prime 5K mouse (10,534 probes, GRCm38.102, 16 nt arms), Prime 5K human (14,862, GRCh38.110), V1 mouse (4,902, GRCm38.102, 20 nt; Mouse Brain gene expression and Mouse Tissue Atlassing merged) and V1 human (16,051, GRCh38.98; Brain, Breast, Colon, Immuno-Oncology, Lung, Multi-Tissue and Cancer, and Skin merged), 46,349 in total. Arm length was taken as half the sum of the BED12 block sizes, the genomic span including the intron. Mouse and human panels were characterized separately within each chemistry. Melting temperature distributions were closely matched between species (Extended Data Fig. 6c) and ligation-dinucleotide usage was concordant across the sixteen dinucleotides (Prime 5K: Spearman rho=0.971; V1: rho=0.973), so a single ruleset per chemistry was used for both.

Two ligation-dinucleotide regimes are apparent. The V1 panels resolve into the three discrete tiers of the published design guidance (https://biotech.illinois.edu/wp-content/uploads/2025/03/Custom-panel-designs.pdf), which the V1 ruleset adopts directly. The Prime 5K panels show a graded preference across sixteen weights, highest for AA and lowest for GC, with the dinucleotides designated optimal in the guidance falling mid-range; the Prime 5K ruleset uses these observed weights. Because the score of an unflagged probe reduces to its dinucleotide weight, median unflagged scores give the weights directly; conditioning on dinucleotide, the median single-flag to unflagged ratio is 0.5 in both chemistries, consistent with the 1/(1+n_flags) form. These weights reflect the design criteria applied to the shipped panels rather than measured ligation efficiency. Prime 5K parameters, the thermal window ([71,77]°C) and the dinucleotide weights, were set with reference to this characterization and to the design scores returned by 10x Genomics for our candidate probes. Xenium V1 parameters (20 nt arms, [68,82]°C) follow the vendor documentation; no fitting was performed for V1. Scores are LIGBASE_XY / (1 + n_flags), rounded to four decimal places, where n_flags counts full-binder Tm outside the chemistry window, a simple tandem repeat, a cytosine run of five or more, and a predicted complete-binder homodimer free energy ≤ -10.5 kcal/mol. Hard filtering is deferred to post-processing so that a candidate pool can be rescored without regeneration. Externally supplied scores, including those returned by 10x Genomics, are retained in separate immutable columns.

### Isoform add-on Panel score comparisons

Score comparisons were performed on the 96-probe isoform add-on panel. Concordance between ProbeHarbour and 10x Genomics design scores was assessed on the 95 add-on probes for which a design score was returned, using Spearman and Pearson correlation and the distribution of absolute differences. Top-ranked ProbeHarbour candidates were compared with the shipped add-on probes at the 93 junctions with an exact same-junction candidate; the three excluded target human-annotated junctions absent from the mouse reference annotation. Differences are given as median and interquartile range. Ligation-dinucleotide concordance between species and between chemistries was computed on the reconstructed predesigned panels as Spearman correlation across the sixteen dinucleotides. Agreement between ProbeHarbour score and observed median QV was assessed on the 96 add-on probes by Spearman correlation and by AUROC at QV≥20.

## Acknowledgments

We thank Dr. Hélène Marie, Dr. Stéphane Martin, and Dr. Bruno Antony for valuable discussions and insightful comments. This study was supported by the Agence Nationale de la Recherche (ANR-25-CE13-7457-01); the European Union’s Horizon Europe programme Marie Skłodowska-Curie Actions Doctoral Network LongTREC (Grant No. 101072892); the France 2030 program (4D-OMICs : ANR-21-ESRE-0052, 3IA : ANR-19-P3IA-0002; France Génomique: ANR-10-INBS-09-03 and 2023-013 X20102; CellID: ANR-24-EXCI-0002); the Canceropole PACA (2025-21Kpole) and the Conseil départemental 06 (2021-367 DGADSH).

## Author contributions

KL conceived and supervised the project. PB, BM, RB and KL secured funding. HC, MI and MP raised the mice; prepared the mouse brains and performed tissue cryo-sectioning and ST slide mounting. MJA performed the Xenium experiments. KL, EM and MF analyzed the data. KL wrote the manuscript. All authors read, updated and approved the final paper.

## Data Availability

All data generated in this study have been deposited in the Gene Expression Omnibus (GEO) and will be made publicly available at the time of publication. External 10x Genomics Xenium, denoted as P56 (Xenium Prime 5k) is accessible at https://www.10xgenomics.com/datasets/xenium-prime-fresh-frozen-mouse-brain. External Oxford Nanopore Technologies Visium HD, denoted as P56M (ONT Visium HD) is accessible at https://epi2me.nanoporetech.com/visium_hd_2025.06/. The isoform add-on panel is browsable at https://probeharbour-96-explorer.streamlit.app/, a ProbeHarbour explorer reporting probeset composition, probe locations on the annotated transcript structures, design scores and empirical per-probe QV from the iSiT experiments.

## Code Availability

Python scripts and codes used for analysis and figures production are freely accessible at https://github.com/cobioda/iSiT. Sparty python package is accessible at https://github.com/cobioda/sparty. ProbeHarbour python package is accessible at https://github.com/cobioda/ProbeHarbour.

## Extended Data Figures

**Extended Data Fig. 1:**
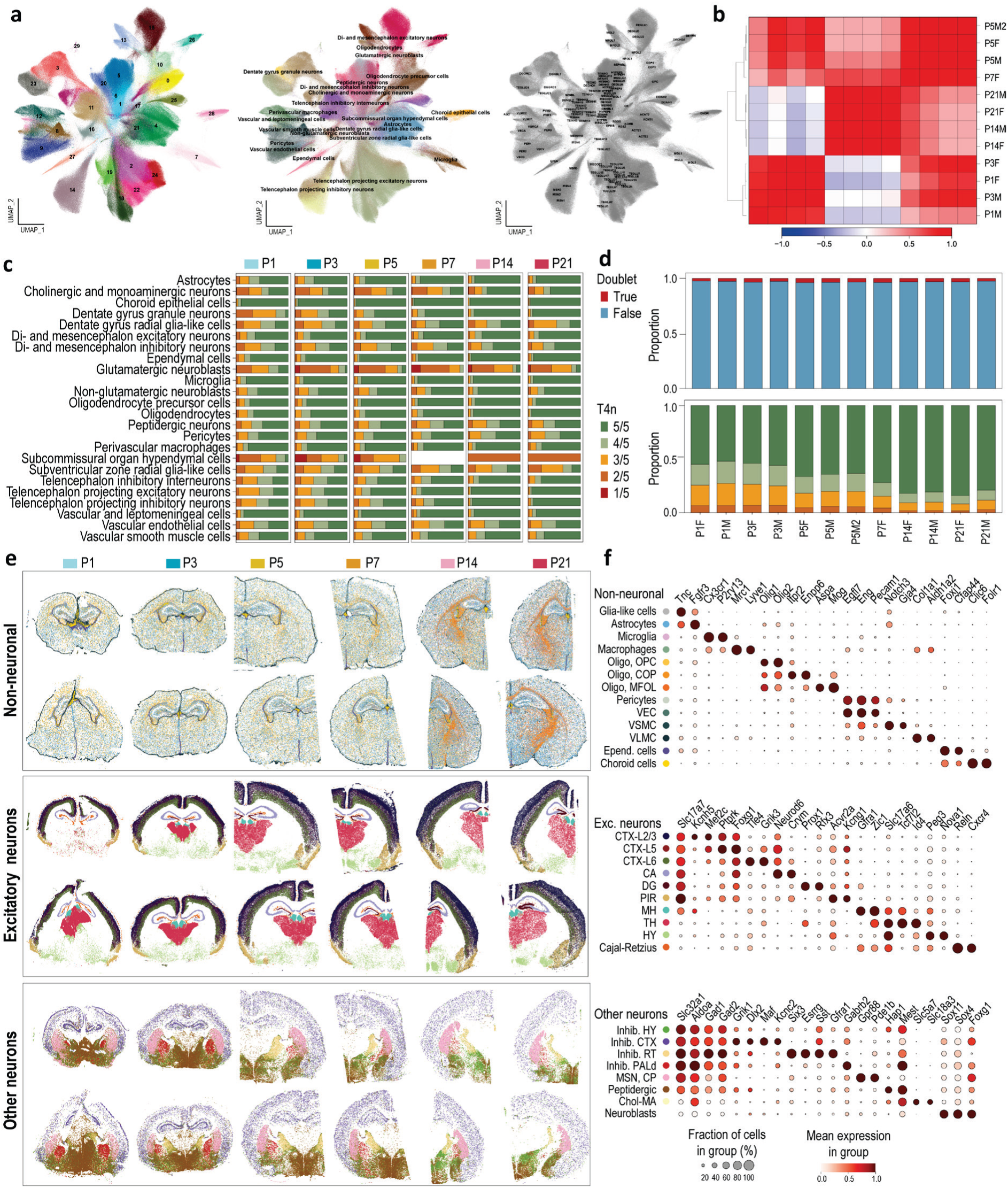
Spatial Transcriptomics cell typing. **a**, Harmony Integration colored by leiden cluster (left), transferred TaxonomyRank4 label (middle) and ClusterName label (right) from Zeisel et al., 2018. **b**, Sample-to-sample correlation plot showing clustering by developmental stage: pup (P1-P3), juvenile (P5-P7) and adolescent (P14-P21). **c**, Robustness of TaxonomyRank4 label transfer, quantified as the number of times the same label is assigned across five independent runs. **d**, Top, proportion of cells classified as doublets per sample, reflecting segmentation inaccuracies. Bottom, proportion of cells per TaxonomyRank4 label transfer robustness, showing improved transfer quality at later stages, consistent with closer similarity to the adolescent reference dataset (Zeisel et al., 2018). **e**, Spatial maps showing non-neuronal cell types (top), excitatory neurons (middle), and inhibitory and other neuronal populations (bottom). **f**, Dot plot of marker gene expression for non-neuronal (top), excitatory neuronal (middle), and other neuronal (bottom) cell types.

**Extended Data Fig. 2:**
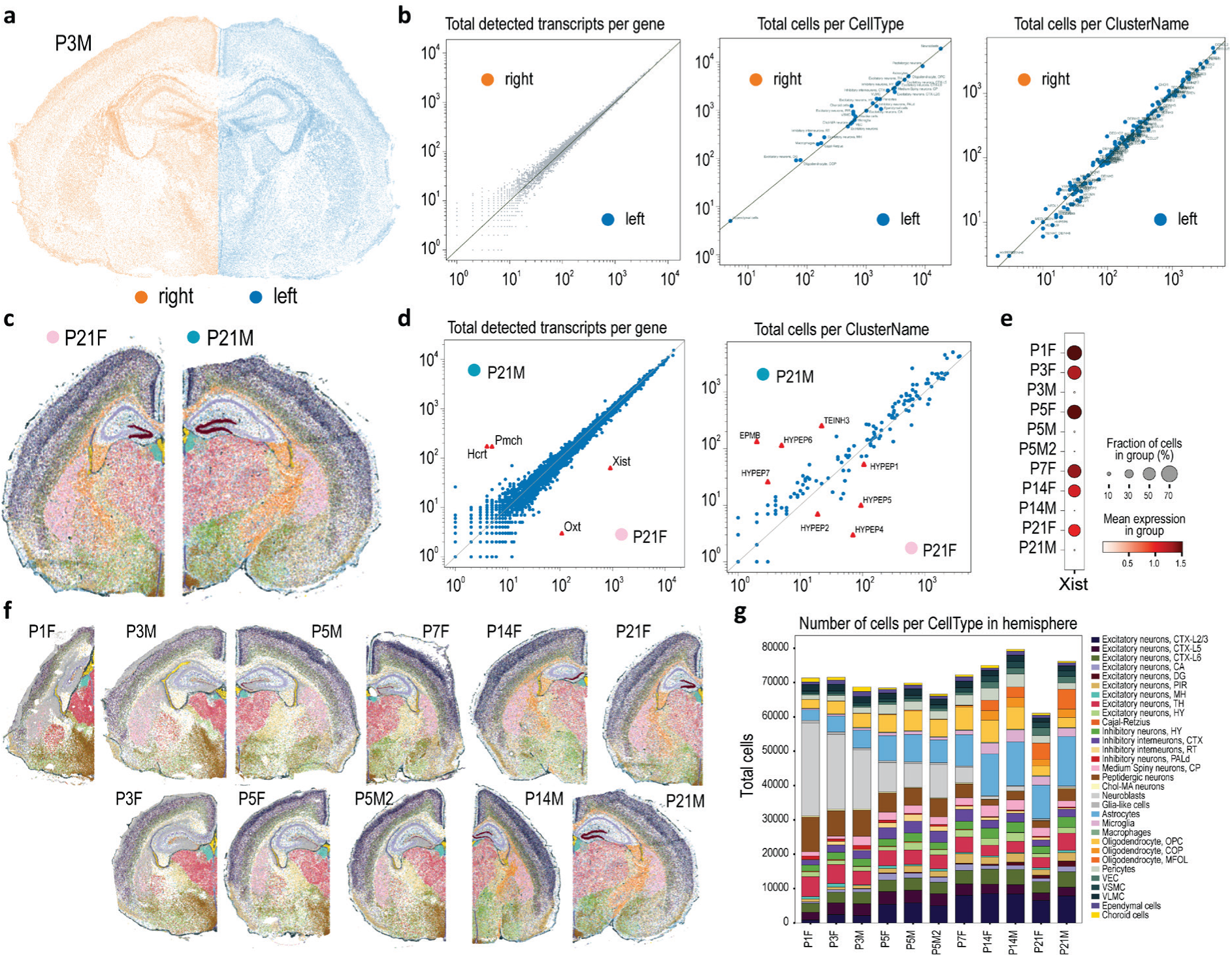
Cell type proportion analysis. **a**, Annotation of left and right hemispheres in the P3M sample. **b**, Comparison between hemispheres showing total detected transcripts per gene (left), total number of cells per cell type (middle), and total number of cells grouped by the Zeisel et al., 2018 ClusterName annotation level (right). **c**, Comparison between male and female samples at the P21 stage. **d**, Total detected transcripts per gene (left) and total number of cells grouped by ClusterName annotation level (right) comparing male and female P21 samples, showing high correlation except for *Xist*, *Oxt*, *Pmch* and *Hcrt*, which are associated with sexually dimorphic hypothalamic neuropeptide neurons. **e**, Mean *Xist* expression per sample. **f**, Visualization of hemisphere extraction across the 11 samples used for cell type proportion comparisons. **g**, Total number of cells per cell type in each hemisphere, per sample.

**Extended Data Fig. 3:**
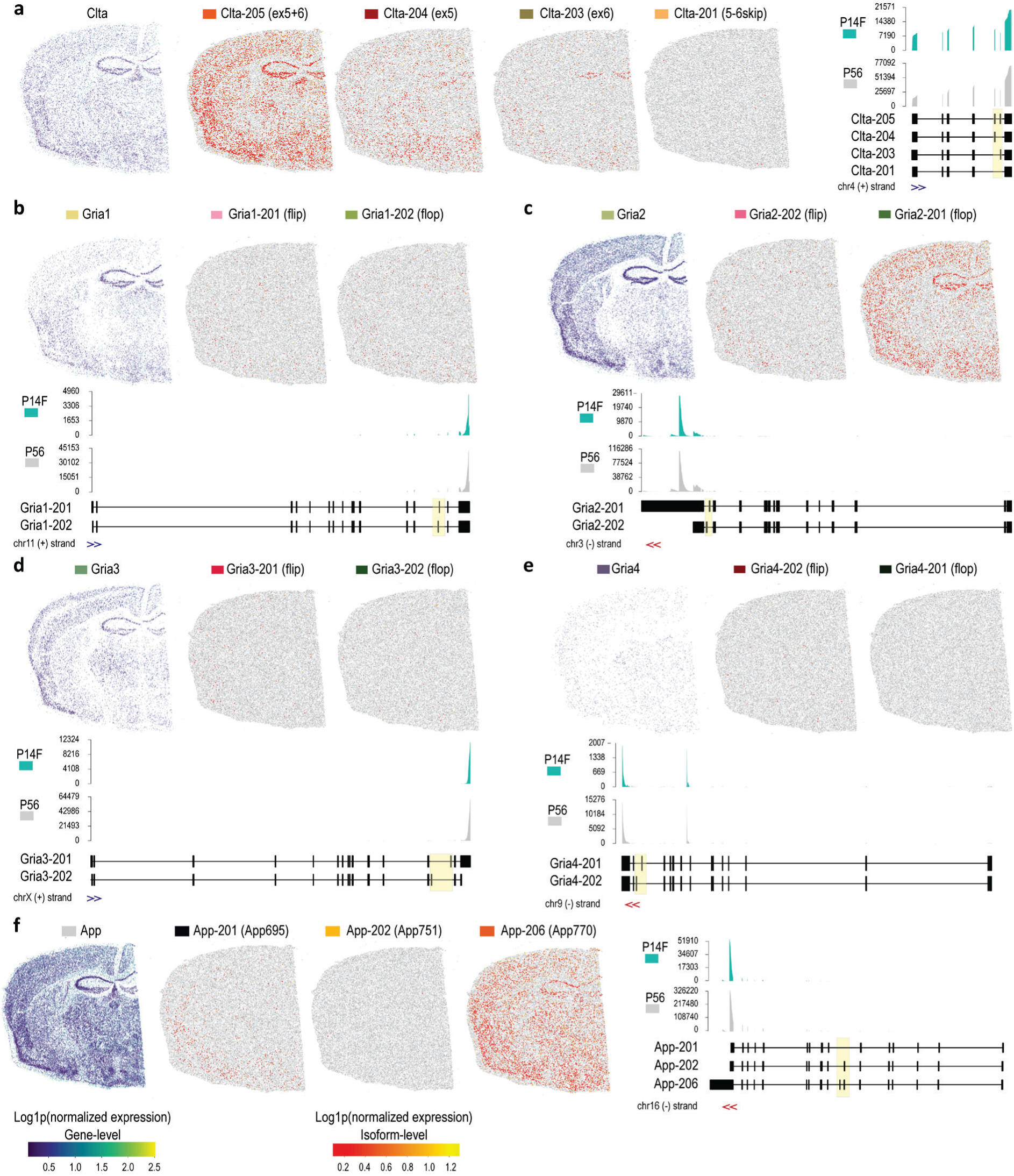
Sequencing-based Spatial isoform Transcriptomics. **a**, P56 expression-level for *Clta* gene-level (left) and Isoform-level (four right panels). Coverage plot is shown for P14F and P56 Visium HD assays. **b**, **c**, **d**, **e**, P56 expression-level for *Gria1*, *Gria2*, *Gria3*, *Gria4* gene-level (left) and Isoform-level (two right panels). Coverage plot is shown for P14F and P56 Visium HD assays. **f**, P56 expression-level for *App* gene-level (left) and Isoform-level (three right panels). Coverage plot is shown for P14F and P56 Visium HD assays.

**Extended Data Fig. 4:**
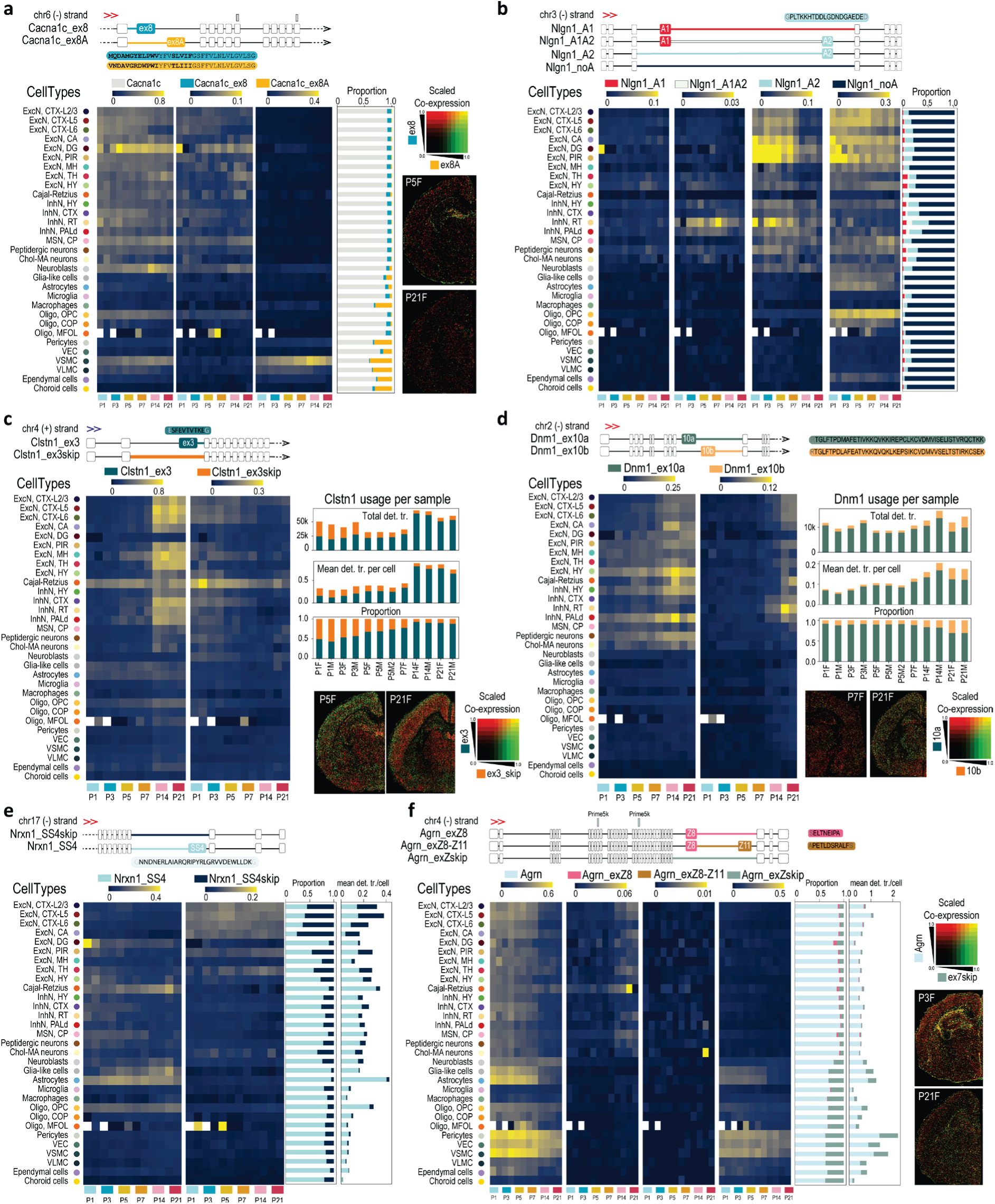
Imaging-based Spatial Isoform Transcriptomics. **a**, *Cacna1c* isoform usage across space and time. **b**, *Nlgn1* isoform usage across space and time. **c**, *Clstn1* isoform usage across space and time. **d**, *Dnm1* isoform usage across space and time. **e**, *Nrxn1* isoform usage across space and time. **f**, *Agrn* isoform usage across space and time.

**Extended Data Fig. 5:**
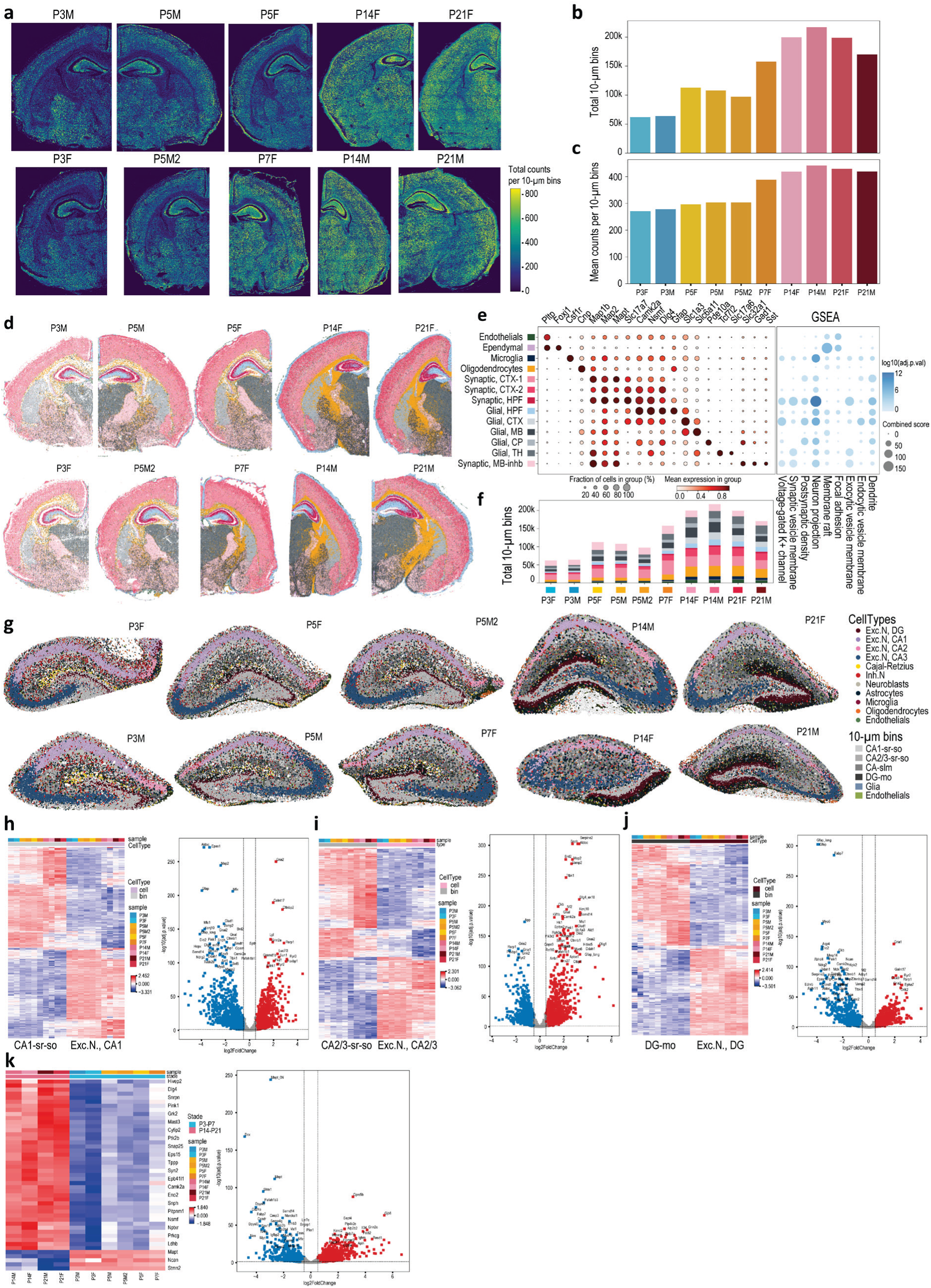
Unassigned RNAs reveal subcellular compartments. **a**, Density of unassigned RNAs per 10-µm bins. **b**, Total number of 10-µm bins per sample, increasing with brain maturation, consistent with the development of cellular protrusions. **c**, Mean transcript counts per 10-µm bins per sample. **d**, Statistical analysis of unassigned RNAs, grouped into 10-µm bins and colored by unsupervised Leiden clustering. **e**, Gene markers for bin clusters and gene set enrichment analysis (GSEA) against the Synaptic Gene Ontology (SynGO) database, identifying clusters associated with dendrites and neuronal projections. **f**, Total number of 10-µm bin per sample colored by bin types as in (e). **g**, Spatial visualization of the hippocampal region for P3, P5, P7, P14 and P21 samples, with cell types and 10-µm bin types overlaid. **h**, **i**, **j**, **k**, Pseudobulk analyses comparing: CA1 pyramidal neurons (CA1-sp) versus CA1 stratum radiatum/stratum oriens bins (CA1-sr-so) (h); CA2/3 pyramidal neurons (CA2/3-sp) versus CA2/3-sr-so bins (i); excitatory neurons in the dentate gyrus (DG) and Ammon’s horn (CA) (DG-sp) versus DG molecular layer bins (DG-mo) (j); and CA1/2/3 sr-so bins between early (P3-P7) and late (P14-P21) developmental stages (k).

**Extended Data Fig. 6:**
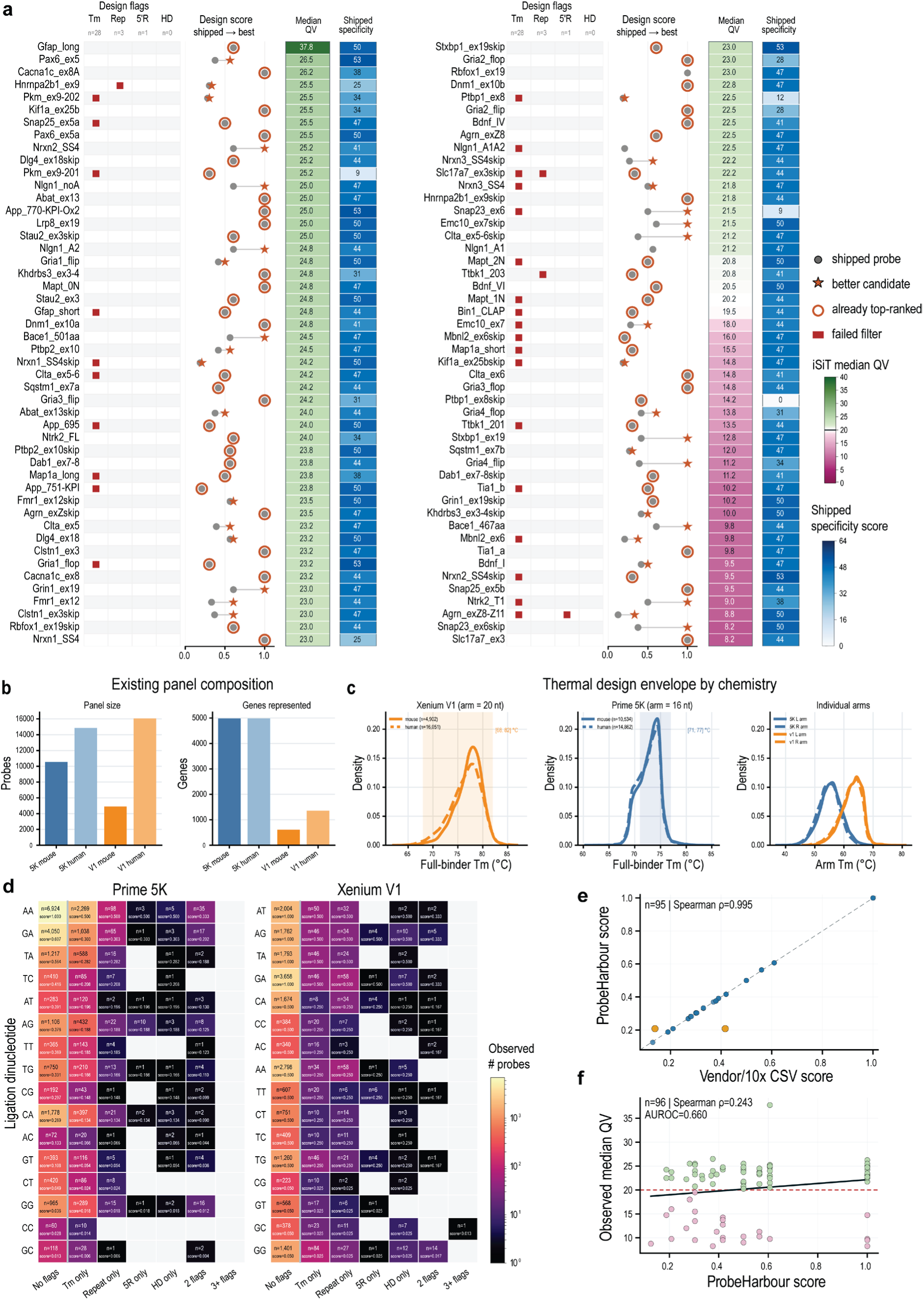
Reverse-engineering of predesigned panels and probe-design validation. **a**, Per-junction re-design over the isoform add-on panel. For each targeted event (rows), the existing-panel-failures block marks shipped probes that fail each design filter (Tm, simple repeat, five-guanine run, homodimer; totals across the panel: Tm n = 28, Rep n = 3, 5’R n = 1, HD n = 0). The design-score block shows the shipped-probe score and the best ProbeHarbour candidate (shipped -> best); markers denote the shipped probe, a better candidate, an already top-ranked probe, and probes failing a filter. Rows are coloured by iSiT median QV (right); 27 of 96 probes fell below QV 20. **b**, Composition of the reverse-engineered panels: number of probes (left) and genes represented (right) for the Prime 5K and V1 mouse and human predesigned panels. **c**, Thermal design envelopes by chemistry. Full-binder Tm density for Xenium V1 (20 nt arms; mouse n = 4,902, human n = 16,051; window [68,82]°C) and Prime 5K (16 nt arms; mouse n = 10,534, human n = 14,862; window [71,77]°C), and per-arm Tm for the left and right arms of each chemistry; dashed lines, human panels; solid lines, mouse panels. **d**, Ligation-dinucleotide preference. Median design score of shipped panel probes under active design rules, resolved by ligation dinucleotide (rows, ordered independently within each chemistry by weight) and by flag category (columns), for Prime 5K and Xenium V1, pooled across species. Cell labels give the median score and probe count; shading gives the number of probes observed. Xenium V1 resolves into three discrete tiers matching the vendor design guidance, whereas Prime 5K shows a graded preference in which a different set of dinucleotides ranks highest. Conditioning on ligation dinucleotide, median scores in each single-flag category are half the corresponding unflagged medians throughout both chemistries, consistent with a 1/(1+n_flags) dependence. **e**, Score concordance: ProbeHarbour score versus the design score returned by 10x Genomics for the add-on panel candidates (n=95; Spearman rho=0.995, Pearson r=0.997; median absolute difference 0.0021; dashed line, y=x). Highlighted points are the two candidates differing by more than 0.01. **f**, Score versus empirical performance: observed median QV versus ProbeHarbour score (n=96; Spearman rho=0.242; AUROC=0.680 for QV≥20; dashed line, QV=20). The score discriminates high-from low-QV probes only weakly, motivating the QV feedback loop in Fig.6a.

